# Metabolism is a major driver of hydrogen isotope fractionation recorded in tree-ring glucose of *Pinus nigra*

**DOI:** 10.1101/2021.07.22.453377

**Authors:** Thomas Wieloch, Michael Grabner, Angela Augusti, Henrik Serk, Ina Ehlers, Jun Yu, Jürgen Schleucher

**Affiliations:** Department of Medical Biochemistry and Biophysics, Umeå University, 901 87 Umeå, Sweden; Institute of Wood Technology and Renewable Materials, University of Natural Resources and Life Sciences Vienna, 3430 Tulln an der Donau, Austria; Research Institute on Terrestrial Ecosystems, National Research Council, 05010 Porano (TR), Italy; Department of Mathematics and Mathematical Statistics, Umeå University, 901 87 Umeå, Sweden

**Keywords:** anaplerotic flux, Calvin-Benson cycle, change point, glucose-6-phosphate shunt, hydrogen stable isotopes, intramolecular isotope analysis, oxidative pentose phosphate pathway, sucrose-to-starch carbon partitioning

## Abstract

– Stable isotope abundances convey valuable information about plant physiological processes and underlying environmental controls. Central gaps in our mechanistic understanding of hydrogen isotope abundances impede their widespread application within the plant and biogeosciences.
– To address these gaps, we analysed intramolecular deuterium abundances in glucose of *Pinus nigra* extracted from an annually resolved tree-ring series (1961 to 1995).
– We found fractionation signals (i.e., temporal variability in deuterium abundance) at glucose H^1^ and H^2^ introduced by closely related metabolic processes. Regression analysis indicates that these signals (and thus metabolism) respond to drought and atmospheric CO_2_ concentration beyond a response change point. They explain ≈60% of the whole-molecule deuterium variability. Altered metabolism is associated with below-average yet not exceptionally low growth.
– We propose the signals are introduced at the leaf-level by changes in sucrose-to-starch carbon partitioning and anaplerotic carbon flux into the Calvin-Benson cycle. In conclusion, metabolism can be the main driver of hydrogen isotope variation in plant glucose.

## Introduction

Stable isotope measurements of the most abundant chemical elements in plants (H, and C) convey valuable information about plant physiological and environmental processes. Plant archives, such as tree rings, preserve this information over millennia enabling elucidation of physiological and environmental dynamics that occur beyond the short timeframes covered by manipulation or monitoring experiments (Peñuelas *et al.*, 2011; Loader *et al.*, 2011; Saurer *et al.*, 2014; Frank *et al.*, 2015; Köhler *et al.*, 2016).

Among all stable isotopes, the hydrogen isotopes protium (^1^H) and deuterium (^2^H, or D) exhibit the largest relative mass difference. As a result, hydrogen isotope effects are normally considerably larger than isotope effects of other elements (Melander & Saunders, 1980). Thus, from the physics point of view, hydrogen isotope compositions of plant compounds (commonly expressed in terms of *δ*D) have a remarkable potential to further our knowledge about plant physiological and environmental processes. However, current plant *δ*D models fail to predict the entire body of available empirical data (see below). Therefore, hydrogen isotopes, in contrast to carbon isotopes, have not yet evolved into standard tools within the plant and biogeosciences.

Current *δ*D models of plant organic matter exhibit three main components: (i) the *δ*D composition of plant water sources, (ii) leaf water D enrichment, and (iii) metabolic fractionations. (i) Plants take up soil water through roots. In most plants, this uptake and water transport through the xylem into leaves occurs without detectable *δ*D shifts (Cernusak *et al.*, 2016; Chen *et al.*, 2020). Additionally, leaf water exchanges with atmospheric water vapour through stomata. Thus, leaf water inherits the *δ*D compositions of both soil water and atmospheric water vapour (Cernusak *et al.*, 2016). (ii) Stomatal evaporation causes D enrichment in leaf water. Adaptations of the Craig-Gordon model describe this process at the site of evaporation (Craig & Gordon, 1965; Flanagan *et al.*, 1991; Farquhar *et al.*, 2007). In a comprehensive series of gas exchange experiments over a range of isotopic and humidity conditions far exceeding natural conditions, Roden and Ehleringer (1999a) confirmed the robustness of a model that approximates leaf water D enrichment (Flanagan *et al.*, 1991). Similarly, on an extensive *δ*D dataset of leaf waters from very dry to very wet field sites, Kahmen *et al.* (2013) confirmed the robustness of the precise *δ*D enrichment model (Farquhar *et al.*, 2007). Reanalysing this latter dataset, Cernusak *et al.* (2016) reported a goodness of fit of *R*^*2*^=0.92 (*n*=332) between modelled and measured *δ*D data. These successful tests for robustness of leaf water *δ*D models strongly suggest that inconsistencies between modelled and measured *δ*D values of plant organic matter as reported by Waterhouse *et al.* (2002) derive from processes that occur downstream of leaf water D enrichment, i.e., from fractionating metabolic processes. (iii) Yakir and DeNiro (1990) developed a *δ*D model describing fractionation in cellulose of aquatic plants and estimated metabolic fractionation factors of autotrophic and heterotrophic growth. To predict *δ*D values of tree-ring cellulose, Roden *et al.* (2000) combined this model with a leaf water *δ*D model (Flanagan *et al.*, 1991) and added a term accounting for *δ*D changes by partial reincorporation of hydrogen from xylem water during cellulose biosynthesis (=heterotrophic hydrogen exchange). Roden and Ehleringer (1999b, 2000) confirmed the robustness of their model for greenhouse- and field-grown riparian trees with good access to water. A robustness test by Waterhouse *et al.* (2002) on tree rings of *Quercus robur* from a dry site failed. These latter authors suggested modelling metabolic fractionations as constants may be inadequate.

All hydrogen positions of leaf-level metabolites inherit D fractionations present in leaf water, the hydrogen source of metabolism. By contrast, metabolic reactions cause D fractionations at specific–intramolecular–hydrogen positions that are directly or indirectly involved in the reaction mechanism (Schmidt *et al.*, 2015). Such fractionations are known to result in pronounced *δ*D differences among hydrogen positions within a metabolite (Martin *et al.*, 1986; Schleucher, 1998; Schmidt *et al.*, 2003). Furthermore, it is now known that metabolic *δ*D variability can occur at a given intramolecular hydrogen position (Schleucher, 1998). Specifically, *δ*D variability at H^6^ of plant leaf glucose was explained by changes of the photorespiration-to-photosynthesis ratio (Schleucher, 1998; Ehlers *et al.*, 2015). Moreover, growing six C_3_ species at ambient CO_2_ concentrations of 280 and 150 ppm, Cormier *et al.* (2018) reported whole-molecule *δ*D differences of ≈20‰ (on average) in leaf α-cellulose due to unknown fractionating metabolic processes. This shows the predictive abilities of plant *δ*D models will improve by accounting for variability in metabolic fractionations. To this end, however, underlying mechanisms need to be elucidated.

Based on findings by Waterhouse *et al.* (2002), we hypothesise metabolic processes manifest significant fractionation signals (throughout the paper, the term ‘signal’ denotes temporal variability in deuterium abundance) at carbon-bound hydrogens in tree-ring glucose under dry conditions. Since metabolic fractionations occur at specific hydrogen positions within molecules, the interpretability of conventional whole-molecule data is fundamentally limited.

For instance, not knowing the intramolecular location of metabolic fractionation impedes attempts to elucidate its enzymatic origin. Additionally, if located at a single hydrogen position, a ≈20‰ whole-molecule effect scales up to ≈140‰ because glucose has seven H-C positions. Therefore, to test our hypothesis, we measured intramolecular D abundances in tree-ring glucose of *Pinus nigra* from a dry site in the Vienna region. This region was chosen for its comprehensive environmental monitoring. We first screen for metabolic fractionation signals. Then, we analyse signal-environment and signal-growth relationships and estimate the contribution of metabolic to whole-molecule fractionation. Lastly, we discuss mechanisms of signal introduction and implications of our findings for isotope studies in the plant and biogeosciences.

## Material and Methods

### Site and samples

*Pinus nigra* Arnold was sampled at Bierhäuselberg, Vienna region, Austria (48.13N, 16.23E, 350 m AMSL). The site exhibits shallow, very dry soil, and an open canopy with apparently low competition among trees (Leal *et al.*, 2008). Sampling targeted dominant trees with umbrella-shaped crowns indicating regular water shortage. From each of 19 specimens (age range: ≈92 to 156 years), two 5 mm stem cores were taken at breast height. Individual tree rings were dated by standard dendrochronological methods (Speer, 2010) and separated using a binocular microscope and a scalpel. To preclude growth-related isotope signals, we analysed tree rings formed from 1961 to 1995 when all trees had reached a comparably stable canopy position. Across trees, all tree-ring material of a given year was combined into annual pools yielding one sample per year. Thus, analytical results represent properties of the tree species at the site rather than individual trees.

### Intramolecular deuterium measurements

After sample randomisation, intramolecular D abundances of glucose were measured following a published protocol (Betson *et al.*, 2006). Samples <20 mg were excluded from analysis (1977, 1978, 1981 and 1982), because too much measurement time would have been required for sufficient precision. Quantitative D NMR spectra (n≥3) were recorded using an AVANCE III 850 with a cryogenic probe optimised for D detection and ^19^F lock (Bruker BioSpin GmbH, Rheinstetten, Germany). Relative intramolecular D abundances were determined by signal deconvolution using Lorentzian line shape fits in TopSpin 4 (Bruker BioSpin GmbH, Rheinstetten, Germany). 3,6-anhydro-1,2-*O*-isopropylidene-α-D-glucofuranose has six methyl-group hydrogens which were introduced during derivatisation and derive from a common batch of acetone. Their D abundance (arithmetic mean) was used as reference to compare glucose D abundances of tree-ring samples from different years as

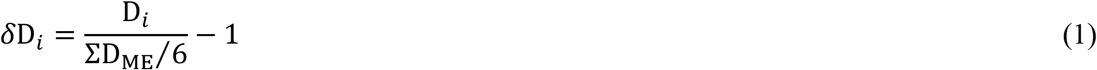

with D_*i*_ and D_ME_ denoting relative D abundances at glucose H-C positions (Fig. 1) and the methyl-group hydrogens, respectively.

### Environmental data

Monthly resolved data of precipitation, *PRE*, air temperature, *TMP*, sunshine duration, *SD*, global radiation, *RAD*, and relative humidity were measured at the climate station Hohe Warte (WMO ID: 1103500). Air vapour pressure deficits, *VPD*, were calculated according to published procedures (Abtew & Melesse, 2013). We used monthly resolved datasets of the self-calibrating Palmer drought severity index, *PDSI*, and the standardised precipitation-evapotranspiration index, *SPEI*, for 48.25N, 16.25E (Wells *et al.*, 2004; Vicente-Serrano *et al.*, 2010). Annual atmospheric CO_2_ concentrations, *C*_a_, were measured at the Mauna Loa observatory, Hawaii by Pieter Tans (NOAA/ESRL, USA) and Ralph Keeling (Scripps Institution of Oceanography, USA). Both the selected grid point and climate station are no more than a horizontal distance of 15 km from the sampling site with a negligible vertical offset. CO_2_ is well mixed in the atmosphere. Thus, all data should represent the site conditions well.

**Figure 1:**
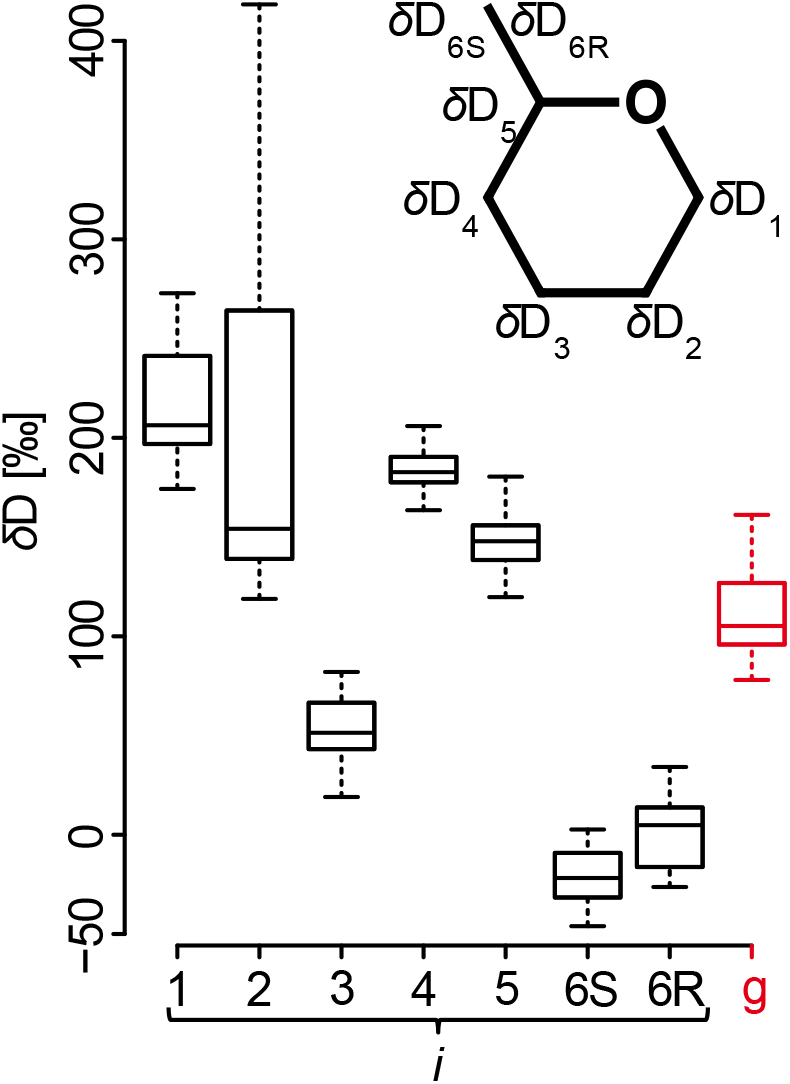
Variability in *δ*D timeseries shown by boxplots. *δ*D_*i*_ and *δ*D_g_ denote timeseries of D abundances at intramolecular H-C positions in tree-ring glucose (black) and of the whole molecule (red), respectively. Data were acquired for tree-ring glucose of *Pinus nigra* laid down from 1961 to 1995 at a site in the Vienna basin (*δ*D_*i*_: ±*SE*=5.4‰, *n*≥3; *δ*D_g_: ±*SE*=3.4‰, *n*≥3). Outliers were removed prior to analysis (*δ*D_1_ to *δ*D_3_: *n*=31, *δ*D_4_ and *δ*D_5_: *n*=30, *δ*D_6S_: *n*=26, *δ*D_6R_: *n*=28, *δ*D_g_: *n*=25). Data reference: Average D abundance of the methyl-group hydrogens of the glucose derivative used for NMR measurements. Insert: Glucose carbon skeleton showing intramolecular locations of *δ*D_*i*_ timeseries.

### Statistical analyses

We calculated the variance contribution of each intramolecular D timeseries, *δ*D_*i*_, to the whole-molecule D timeseries, *δ*D_g_, as follows. With

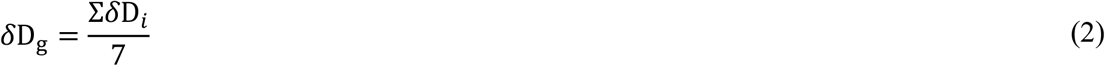

the variance of *δ*D_g_ equals the covariance average between *δ*D_*i*_ and *δ*D_g_ as

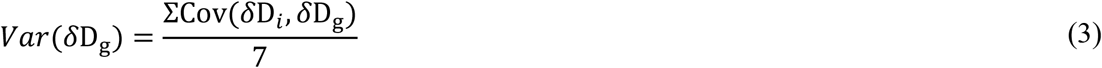

Thus, variability contributions of *δ*D_*i*_ to *δ*D_g_ are given as

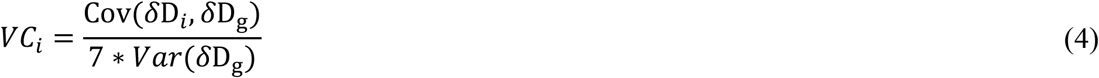

See Notes S1 for information on other statistical analyses.

## Results

To better understand metabolic fractionations, we measured intramolecular D abundances at all seven H-C positions of glucose extracted across an annually resolved *Pinus nigra* tree-ring series (Fig. 1). Our timeseries covers the period 1961 to 1995 but lacks data for 1977, 1978, 1981 and 1982. Hence, the dataset consists of seven intramolecular *δ*D_*i*_ timeseries each comprising 31 observations (*n*=7*31=217). Additionally, we derived the molecular average timeseries, *δ*D_g_ (*n*=31).

### Outlier analysis

Outliers can potentially impair statistical analyses because they may reflect either experimental errors or extreme states of the system. Since we were interested in the normal functioning of plant metabolism and metabolic fractionation, we removed outliers to ensure robust statistical results. We identified outliers in *δ*D_*i*_ timeseries (n=31) as values that were 1.5 times the interquartile range below the first or above the third quartile. We removed one data point each from *δ*D_4_ and *δ*D_5_ (*n*=30), three data points from *δ*D_6R_ (*n*=28) and five data points from *δ*D_6S_ (*n*=26) resulting in six missing data points in *δ*D_g_ (*n*=25). Average standard errors of *δ*D_*i*_ and *δ*D_g_ measurements are 5.4‰ and 3.4‰, respectively.

### *δ*D_1_ and *δ*D_2_ reflect highly variable and closely related metabolic processes

Intramolecular D abundances of *Pinus nigra* tree-ring glucose differ among H-C positions (Figs. 1, 2b, and Notes S2). Medial intramolecular differences exceed 225‰ (*δ*D_1_ vs. *δ*D_6S_). This shows that metabolic fractionations, being position-specific, have strong effects. Interestingly, in some years, intramolecular *δ*D differences approach 450‰ (Notes S2). Thus, metabolic fractionations are highly variable on the interannual scale.

**Figure 2:**
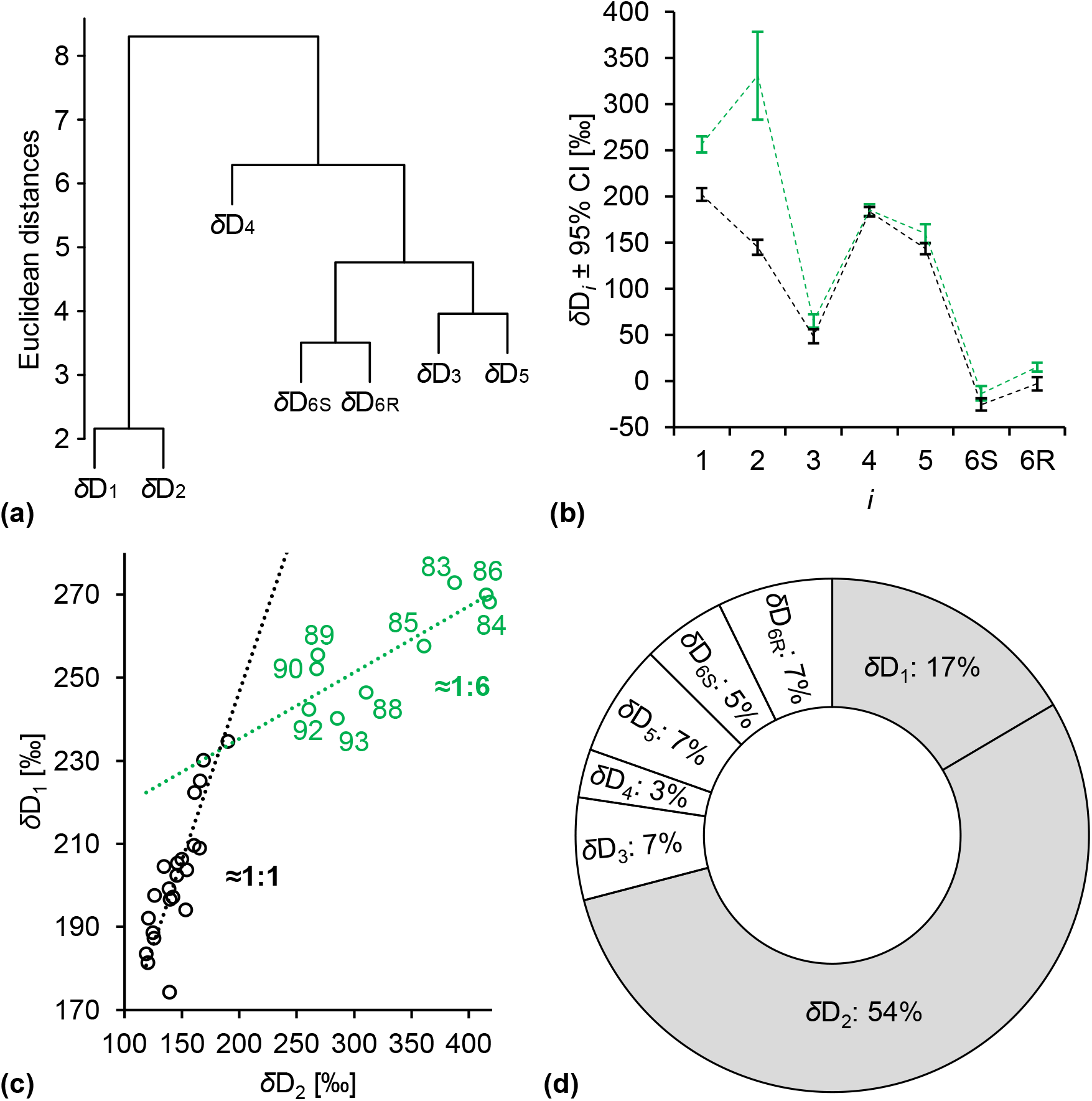
**(a)** Common variability among *δ*D_*i*_ timeseries detected by Hierarchical Cluster Analysis (HCA). **(b)** Intramolecular *δ*D_*i*_ patterns of the low- and high-value group (black and green, respectively). **(c)** Relationship between *δ*D_1_ and *δ*D_2_ (overall *R*^*2*^=0.93, *p*<10^−15^, *n*=31). Dotted lines: Linear major axis regression models showing that *δ*D_1_ and *δ*D_2_ increase at a 1:1 ratio at low values (black) and at a 1:6 ratio at high values (green). **(d)** Percent contributions of *δ*D_*i*_ to the variance in *δ*D_g_. *δ*D_*i*_ and *δ*D_g_ denote timeseries of D abundances at intramolecular H-C positions in tree-ring glucose and of the whole molecule, respectively. Data were acquired for tree-ring glucose of *Pinus nigra* laid down from 1961 to 1995 at a site in the Vienna basin (*δ*D_*i*_: ±*SE*=5.4‰, *n*≥3; *δ*D_g_: ±*SE*=3.4‰, *n*≥3). Outliers were removed prior to analysis (*δ*D_1_ to *δ*D_3_: *n*=31, *δ*D_4_ and *δ*D_5_: *n*=30, *δ*D_6S_: *n*=26, *δ*D_6R_: *n*=28, *δ*D_g_: *n*=25). Low- and high-value group identified by HCA (Notes S2). Figure 2b shows discrete data. Dashed lines added to guide the eye. The variance partitioning analysis (2d) is based on years without missing data after removing outliers (*n*=8*25). Data reference: Average D abundance of the methyl-group hydrogens of the glucose derivative used for NMR measurements.

Hierarchical cluster analysis (HCA) groups timeseries according to co-variability. Timeseries carrying common signals, i.e., information about a common process or strongly related processes, form clusters. Performing HCA on our *δ*D_*i*_ dataset, we found a strong separation between a cluster comprising *δ*D_1_ and *δ*D_2_ and a cluster comprising *δ*D_3_ to *δ*D_6R_ (Fig. 2a). Additionally, we found separations within the *δ*D_3_ to *δ*D_6R_ cluster, with *δ*D_4_ showing the highest degree of independency. Thus, *δ*D_*i*_ timeseries carry information about several fractionation processes.

While *δ*D_3_ to *δ*D_6R_ exhibit similar degrees of variability (Fig. 1; *SD*=±10.5‰ to ±16.8‰, range=42‰ to 63‰, *n*=26 to 31), *δ*D_2_ varies considerably (*SD*=±91.8‰, *range*=300‰, *n*=31) and *δ*D_1_ is also relatively variable (*SD*=±28.4‰, *range*=99‰, *n*=31). Increased D variability at two out of seven H-C positions indicate highly variable metabolic fractionations at these positions, H^1^, and H^2^.

To further investigate this, we performed HCA on annual *δ*D_*i*_ patterns and found two groups (Notes S2). Figure 2b shows the arithmetic average patterns of both groups. While *δ*D_3_ to *δ*D_6R_ values remain comparably constant under all conditions experienced by the trees, *δ*D_1_ and *δ*D_2_ separate annual *δ*D_*i*_ patterns into a low-value group (black, *n*=22) and a high-value group (green, *n*=9). Average and median increases in *δ*D_1_ and *δ*D_2_ are statistically significant (one-tailed Student’s t-tests: *δ*D_1_, *p*<10^−9^; *δ*D_2_, *p*<10^−4^; one-tailed Mood’s median tests: *δ*D_1_ and *δ*D_2_, *p*<.001). Since differences in *δ*D_*i*_ patterns reflect differences in metabolism, this finding confirms that glucose H^1^ and H^2^ carry exceptionally variable metabolic fractionation signals. Underlying metabolic processes are closely related (Fig. 2a).

Non-metabolic fractionation processes (e.g., leaf water D enrichment) have equal effects on all *δ*D_*i*_. In regression analysis investigating relationships among different *δ*D_*i*_, these processes can be expected to yield slopes=1. By contrast, metabolic fractionation processes affect subsets of *δ*D_*i*_ and may thus cause deviations from slope=1. To investigate which kind of fractionation shapes the variation in both the low- and high-value group, we performed major axis regression analysis on low and high *δ*D_1_ and *δ*D_2_ values (Fig. 2c, black and green circles, respectively). Compared to ordinary least squares regression analysis, major axis regression analysis yields truer parameter estimates when x-variable data contain significant relative errors as is the case here for low-value data. For low values, we found a slope of ≈0.81 with a 95% confidence interval of ≈0.60 to ≈1.07 (*n*=22), i.e., *δ*D_1_ and *δ*D_2_ increase at a ≈1:1 ratio. For high values, we found a slope of ≈0.16 with a 95% confidence interval of ≈0.07 to ≈0.25 (*n*=9), i.e., *δ*D_1_ and *δ*D_2_ increase at a ≈1:6 ratio. Thus, while *δ*D_1_ and *δ*D_2_ variability in the low-value group is consistent with expectations related to non-metabolic fractionation processes, high *δ*D_1_ and *δ*D_2_ values reflect additional effects by metabolic fractionation processes.

### Metabolism changes in response to dry conditions after a change point is crossed

To investigate the controls of the metabolic fractionation processes, we estimated metabolic fractionation at glucose H^1^ and H^2^ as

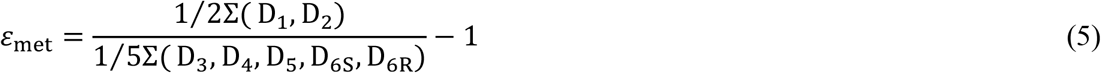

where D_*i*_ denotes relative D abundances at glucose H-C positions (Fig. 1). This procedure maximises metabolic fractionation signals located at H^1^ and H^2^, removes fractionation signals equally present at all H-C positions (leaf water D enrichment), and minimises signals of processes affecting all H-C positions similarly (heterotrophic hydrogen exchange). Weaker metabolic fractionation signals present at glucose H^3^ to H^6^ (Figs. 1, and 2a) will be further explored in future analyses of the dataset.

In general, the fractionating metabolic processes may be under developmental control (Gray & Song, 1984). However, formation of tree rings analysed here occurred when the trees had reached a stable canopy position. On average, tree-ring width measurements of the samples reach back to 1865 (*range*: 1840 to 1918) and *δ*D_*i*_ timeseries start at 1961. Thus, the fractionating metabolic processes can be expected to be controlled by environmental parameters. In the following, these are elucidated in six steps which motivate two integrated isotope-environment models presented subsequently.

#### Step 1

Increases of *ε*_met_ are temporally confined to the second part of the study period (Fig. 2c, 1983 to 1993). This may indicate the presence of a response change point. To test this, we performed a batch change point analysis based on the non-parametric Mann-Whitney-Wilcoxon test (Ross, 2015). We found a change point at 1980 (*p*<.001, *n*=31) showing that the observed frequency distribution of *ε*_met_ (Notes S3) does not align with the properties of a single theoretical probability distribution. That is, *ε*_met_ data of 1961 to 1980 and 1983 to 1995 follow different probability distributions (1981 and 1982 are missing). Thus, the *ε*_met_ timeseries exhibits a change point beyond which the fractionating metabolic processes are activated or upregulated.

#### Step 2

Upregulation of the fractionating metabolic processes occurs beyond a change point. Before the change point, environmental variability exerts no significant control (Fig. 2c, 1:1 ratio). To investigate when the change point is crossed, we correlated *ε*_met_ with different climate parameters. Since *ε*_met_ does not follow a normal distribution (Notes S3), we calculated Spearman’s rank correlation coefficient, *ρ*. Our analysis included four time periods: January to December, year; March to May, spring; March to November, growing season (Wieloch *et al.*, 2018); and June to July, summer. For most periods, we found significant positive correlations of *ε*_met_ with *VPD* and significant negative correlations with *PDSI* and *SPEI* (Table 1). High *VPD*, low *PDSI*, and low *SPEI* correspond to dry conditions. Thus, the results indicate that upregulation of the fractionating metabolic processes (high *ε*_met_) occurs in response to dry conditions.

**Table 1:**
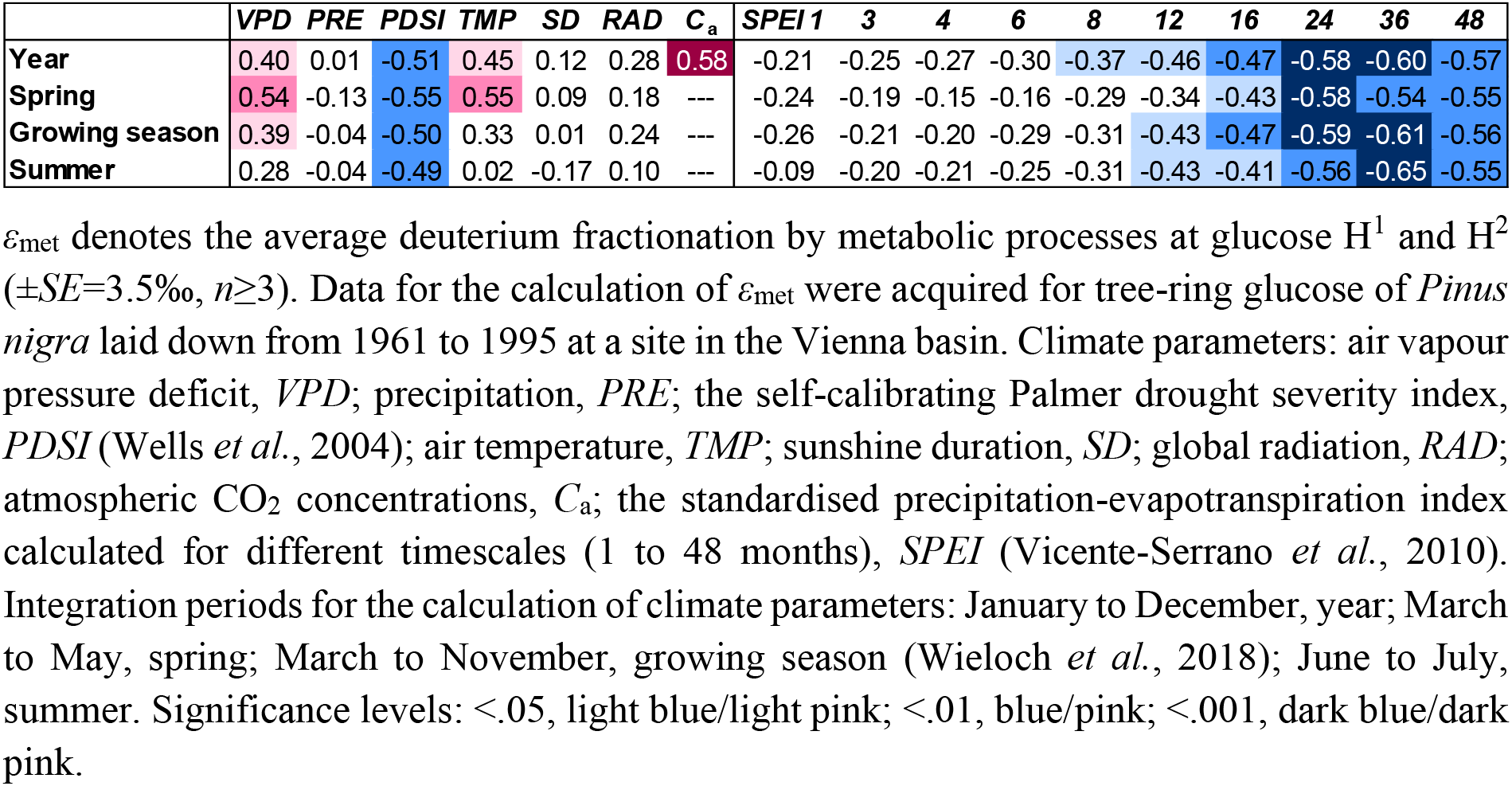
Spearman’s rank correlation coefficients and significance levels for relationships between *ε*_met_ and climate parameters within the period 1961 to 1995 (*n*=31).

#### Step 3

*SPEI* can be calculated for different timescales indicating different types of hydrological drought (Vicente-Serrano *et al.*, 2010). While short timescales indicate variability in soil moisture, long timescales indicate variability in groundwater storage (Vicente-Serrano *et al.*, 2010). To investigate effects of different drought types on the fractionating metabolic processes, we correlated *ε*_met_ with *SPEI* calculated for timescales of 1, 3, 4, 6, 8, 12, 16, 24, 36, and 48 months. We found increasing correlation coefficients from short to long timescales (Table 1) indicating that long-term drought promoted upregulations of the fractionating metabolic processes (possibly by groundwater depletion). Long-term drought events have a low temporal frequency (Vicente-Serrano *et al.*, 2010). This may explain why upregulations of the fractionating metabolic processes occurs not scattered over the entire study period but over consecutive years during the second half of the study period (Fig. 2c).

#### Step 4

In addition to correlations indicative of drought control, we found a significant positive correlation between *ε*_met_ and *C*_a_ (Table 1) which may constitute a pseudocorrelation resulting from intercorrelation between *C*_a_ and long-term drought (Pearson correlation between *C*_a_ and the growing season 48-months *SPEI*: *r*=-0.83, *p*<10^−7^, *n*=31). For the year and spring, we found significant positive correlations between *ε*_met_ and *TMP* (Table 1). Thus, air temperature may contribute to upregulations of the fractionating metabolic processes. Alternatively, the *TMP* correlation may be explained by intercorrelation with drought parameters. Overall, long-term drought shows the highest correlation coefficients (Table 1) indicating that it is the most important environmental control. Furthermore, we considered pathogens as cause for upregulations of the fractionating metabolic processes. The Research Centre for Forests (Vienna, Austria) closely monitors and reports on pathogen status. To our knowledge, there are no reports of major damage by pathogens for the studied *Pinus nigra* stand during the 1980’s and early 1990’s. Thus, significant effects of pathogens on upregulations of the fractionating metabolic processes are unlikely. Similarly, effects by forest management interventions are unlikely since the stand is not used economically (Leal *et al.*, 2008). Effects by air pollutants could not be assessed due to limited data availability.

#### Step 5

To assess the magnitude of the drought shift indicated by correlation, we plotted the growing-season 48-months *SPEI* as function of time (Fig. 3a). Trendline analysis shows a significant increase in drought severity with a long-term shift from ‘non-drought’ from 1961 to 1978, to ‘mild drought’ from 1979 to 1988, to ‘moderate drought’ from 1989 to 1995 according to the classification of drought severity (*R*^*2*^=0.69, *p*<10^−9^, *n*=35) (McKee *et al.*, 1993; Vicente-Serrano *et al.*, 2010). Thus, upregulations of the fractionating metabolic processes in between 1983 and 1993 occurred when the study region experienced mild to moderate drought. Note that actual conditions at the site may have deviated from regional conditions plotted in figure 3a.

**Figure 3:**
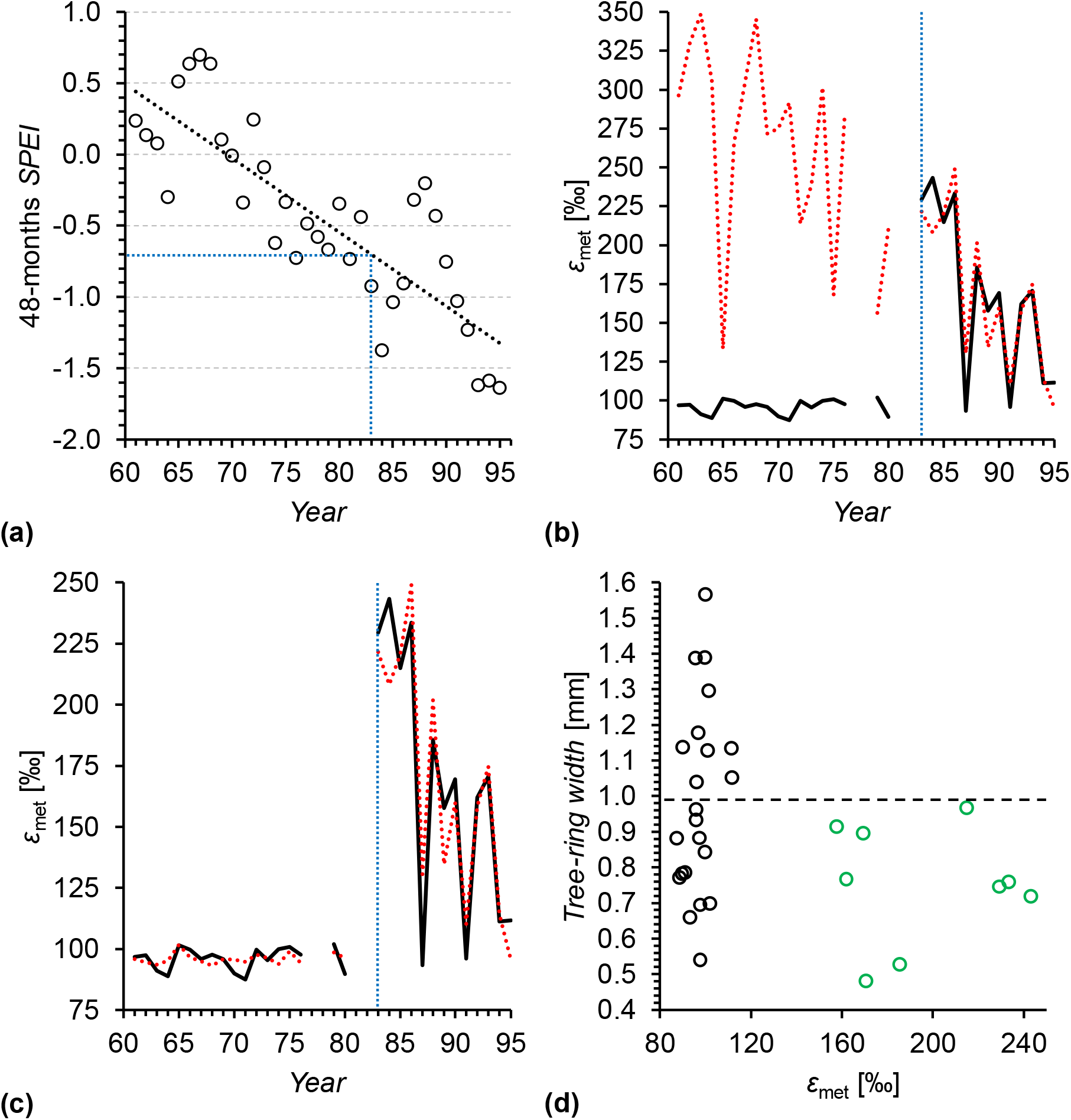
**(a)** Long-term development of growing season drought in the study region. Trendline analysis shows a significant increase in drought severity with shifts from “non-drought” from 1961 to 1978, to “mild drought” from 1979 to 1988, to “moderate drought” from 1989 to 1995 (dotted black line, *R*^*2*^=0.69, *p*<10^−9^, *n*=35). *SPEI* denotes the standardised precipitation-evapotranspiration index (Vicente-Serrano *et al.*, 2010). Here, it was calculated for the growing season (March to November (Wieloch *et al.*, 2018)) at 48.25° N, 16.25° E on a 48-months timescale indicative of long-term drought variability (black circles). Expressions of *SPEI* values indicate the drought severity as ≥-0.49, non-drought; −0.5 to −0.99, mild drought; −1 to −1.49, moderate drought; −1.5 to −1.99, severe drought; <-2, extreme drought (McKee *et al.*, 1993; Vicente-Serrano *et al.*, 2010). **(b) and (c)** Comparison of measured (solid black line) and modelled (dotted red line) metabolic fractionation at glucose H^1^ and H^2^, *ε*_met_ (*n*=31). In (b) and (c), data were modelled by a multivariate linear model (Eq. 6) and a multivariate change point model (Eq. 7, Table 3) (Fong *et al.*, 2017), respectively. Dotted blue line: Response change point as determined by the change point model. **(d)** Relationship between *ε*_met_ and *tree-ring width*. Green and black circles: Years in which the fractionating metabolic processes were upregulated and downregulated, respectively (Fig. 2). Dashed line: Average tree-ring width of years in which the fractionating metabolic processes were downregulated (black circles, 0.99 mm). Tree-ring widths were measured for previous dendro-ecological studies (Leal *et al.*, 2008). Data for the calculation of *ε*_met_ were acquired for tree-ring glucose of *Pinus nigra* laid down from 1961 to 1995 at a site in the Vienna basin (±*SE*=3.5‰, *n*≥3).

**Table 2:**
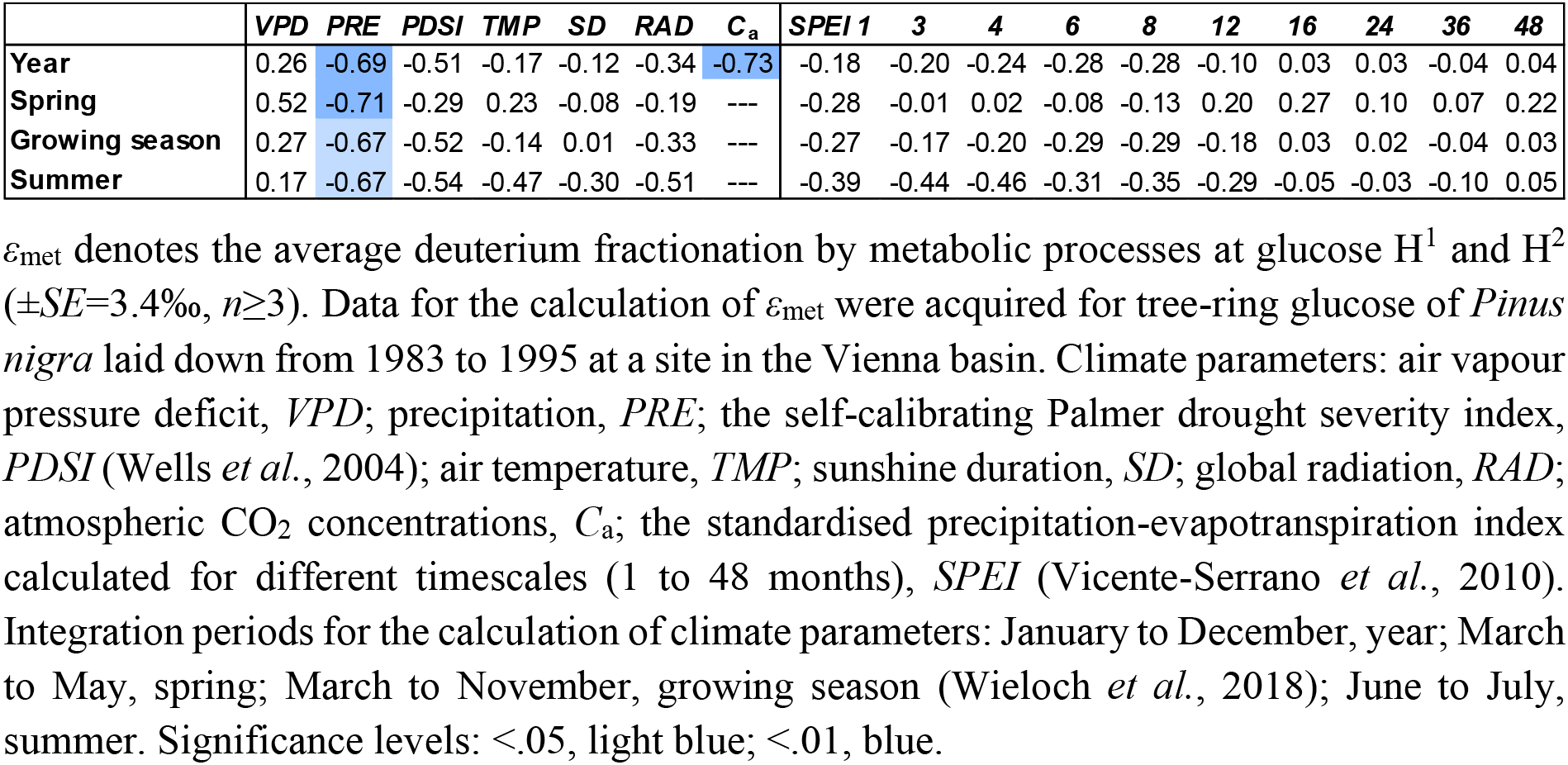
Pearson correlation coefficients and significance levels for relationships between *ε*_met_and climate parameters within the period 1983 to 1995 (*n*=13).

**Table 3:**
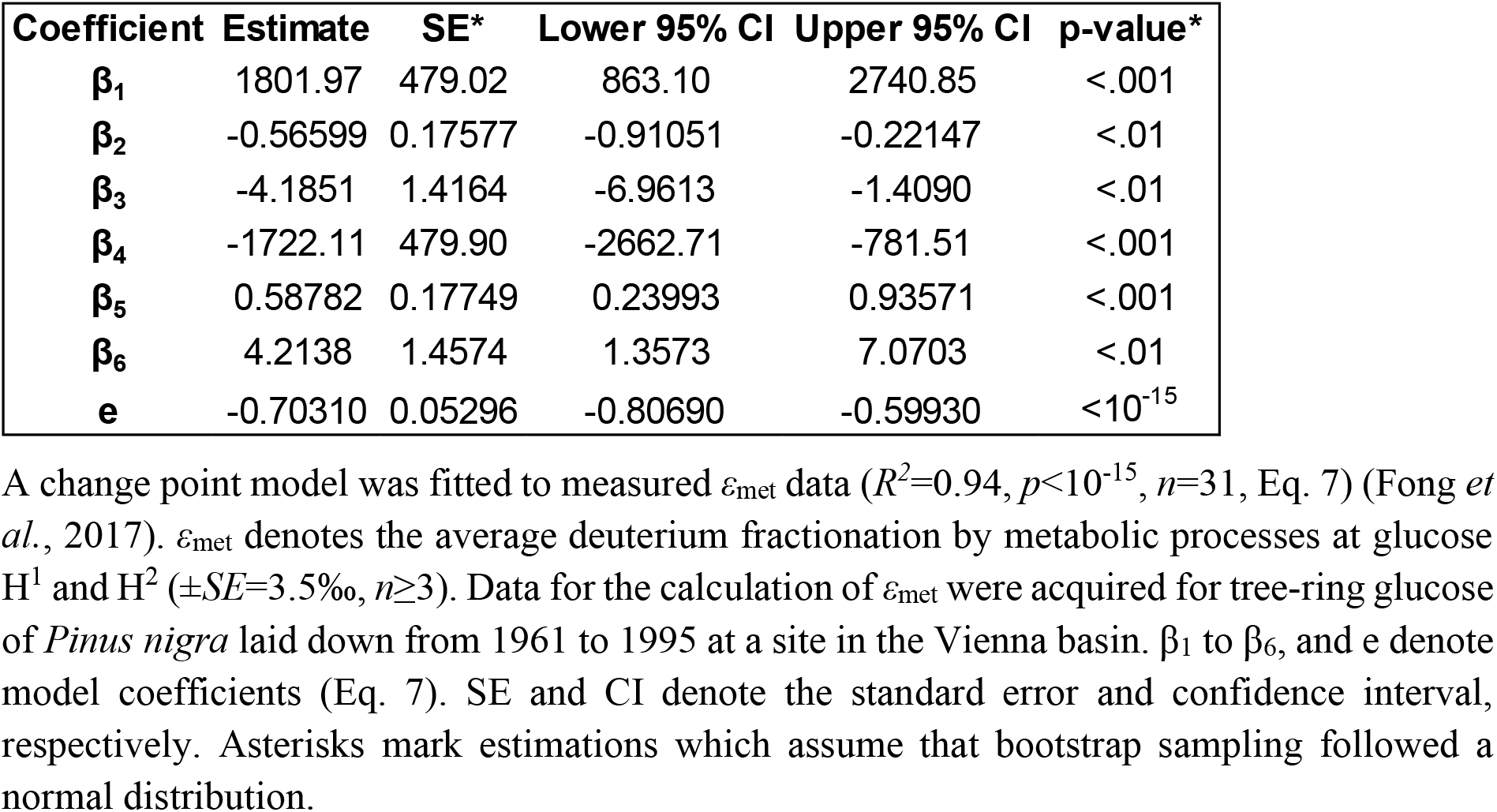
Estimated coefficients of the *ε*_met_ change point model.

| Coefficient | Estimate | SE* | Lower 95% CI | Upper 95% CI | p-value* |
| --- | --- | --- | --- | --- | --- |
| $\beta_1$ | 1801.97 | 479.02 | 863.10 | 2740.85 | <.001 |
| $\beta_2$ | -0.56599 | 0.17577 | -0.91051 | -0.22147 | <.01 |
| $\beta_3$ | -4.1851 | 1.4164 | -6.9613 | -1.4090 | <.01 |
| $\beta_4$ | -1722.11 | 479.90 | -2662.71 | -781.51 | <.001 |
| $\beta_5$ | 0.58782 | 0.17749 | 0.23993 | 0.93571 | <.001 |
| $\beta_6$ | 4.2138 | 1.4574 | 1.3573 | 7.0703 | <.01 |
| e | -0.70310 | 0.05296 | -0.80690 | -0.59930 | <10<sup>-15</sup> |

A change point model was fitted to measured $\varepsilon_{\text{met}}$ data ( $R^2=0.94$ , $p<10^{-15}$ , $n=31$ , Eq. 7) (Fong *et al.*, 2017). $\varepsilon_{\text{met}}$ denotes the average deuterium fractionation by metabolic processes at glucose $\text{H}^1$ and $\text{H}^2$ ( $\pm\text{SE}=3.5\text{‰}$ , $n\geq 3$ ). Data for the calculation of $\varepsilon_{\text{met}}$ were acquired for tree-ring glucose of *Pinus nigra* laid down from 1961 to 1995 at a site in the Vienna basin. $\beta_1$ to $\beta_6$ , and e denote model coefficients (Eq. 7). SE and CI denote the standard error and confidence interval, respectively. Asterisks mark estimations which assume that bootstrap sampling followed a normal distribution.

#### Step 6

In 1994 and 1995, long-term drought was exceptionally strong (Fig. 3a). However, this did not result in significant upregulations of the fractionating metabolic processes (Fig. 2c). Therefore, we hypothesise long-term drought beyond a critical level is merely setting the stage for upregulation and secondary climate factors exert control on shorter timescales. To investigate this, we repeated the *ε*_met_-climate-correlation analysis with data of the second half of the study period, 1983 to 1995. We justify this data selection as follows. In 1983, upregulation of the fractionating metabolic processes occurred for the first time (Fig. 2c). From 1983 to 1995, long-term drought became more severe (Fig. 3a, dotted black line). Thus, the site conditions during this period may have been generally favourable for upregulation. Before 1983, long-term drought was less severe and may have rendered upregulations impossible (Fig. 3a). In this scenario, control by secondary climate factors would have been impossible and considering their variability would impair correlation results. Since the selected subset of *ε*_met_ (1983 to 1995) follows a normal distribution (Notes S3), we calculated Pearson correlation coefficients, *r*. We found significant negative correlations between *ε*_met_ and *PRE* (Table 2). Under generally dry conditions, low precipitation may result in low soil moisture. Thus, the results indicate that upregulations of the fractionating metabolic processes (high *ε*_met_) occurred in response to short-term depletions in soil moisture when long-term drought was beyond the critical level. In this respect, it is noteworthy that *Pinus nigra* reportedly has a well-developed lateral root system in upper soil layers to access soil water but can additionally tap into deep water sources if available (Stokes *et al.*, 2002; Peñuelas & Filella, 2003). Furthermore, we found a significant negative correlation between *ε*_met_ and annual *C*_a_ (Table 2). This correlation is not explained by intercorrelation between *C*_a_ and long-term drought (Pearson correlation between *C*_a_ and the growing season 48-months *SPEI*: *r*=-0.39, *p*>.1, *n*=13). Thus, the result indicates that high atmospheric CO_2_ concentrations counteracted upregulations of the fractionating metabolic processes (high *ε*_met_) when long-term drought was beyond the critical level.

#### Model 1

To further corroborate the existence of a response change point (Step 1), and the requirement of long-term drought (Steps 2, and 3) for the manifestation of effects by secondary climate factors (Step 6), we modelled two *ε*_met_ groups as functions of *PRE* and *C*_a_ as

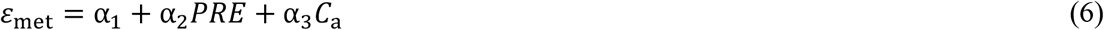

While the first group includes data of 1961 to 1982 corresponding to less severe long-term drought, the second group includes data of 1983 to 1995 corresponding to more severe long-term drought (Fig. 3a). For the latter group, we found adequate linear models including, as explanatory variables, spring *PRE* and annual *C*_a_ (*R*^*2*^=0.74, *p*<.01, *n*=13), and summer *PRE* and annual *C*_a_ (*R*^*2*^=0.78, *p*<.001, *n*=13). The most adequate model we found includes summed spring and summer *PRE* and annual *C*_a_ (*R*^*2*^=0.87, *p*<10^−4^, *n*=13, α_1_=1801.97, α_2_=-0.566, α_3_=4.1851; see Notes S4 for bivariate relationships of *ε*_met_ with *PRE* and *C*_a_, respectively). Both *PRE* and *C*_a_ contributed significantly to all models (*p*<.05). For *ε*_met_ data of the first group, 1961 to 1982, the most adequate model has no explanatory power (*p*>.1, *n*=18). Modelling *ε*_met_ by the most adequate model works well for the second group, 1983 to 1995, yet produces large offsets between all measured and modelled values of the first group (Fig. 3b). This corroborates that (i) a response change point exists, and (ii) the manifestation of effects by secondary climate factors requires long-term drought. Furthermore, it should be noted that the most adequate model accounts well for low *ε*_met_ values in 1994 and 1995 (Fig. 3b). During these years, long-term drought was exceptionally severe, yet *PRE* and *C*_a_ were high (Figs. 3a, Notes S4). Thus, long-term drought alone may not necessarily lead to upregulation of the fractionating metabolic processes. High precipitation during spring and summer as well as high atmospheric CO_2_ concentrations may exert counteracting effects.

#### Model 2

Above, we found indications for several properties of *ε*_met_ of *Pinus nigra* tree-ring glucose. To test these properties in an integrated way and to estimate the change point value, we fitted a change point model to the entire dataset (Fong *et al.*, 2017). It takes the form

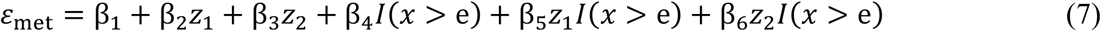

where β_1_ to β_6_ are model coefficients, *z*_1_ is March to June *PRE*, *z*_2_ is annual *C*_a_, *x* is the trend of the growing season 48-months *SPEI* used as change point variable (Fig. 3a, dotted black line), and e is the change point value. If *x*>e then *I*(*x*>e) equals 1, else *I*(*x*>e) equals 0. The fitted model explains 94% of the variability in *ε*_met_ and is highly significant (*p*<10^−15^, *n*=31). All explanatory variables and the change point contribute significantly (Table 3). The estimated change point is at −0.7 ± 0.05SE corresponding to mild long-term drought (*p*<10^−15^, Table 3, Fig. 3a). The presence/absence and directionality of modelled *ε*_met_ relationships with precipitation and atmospheric CO_2_ concentration agree with findings given above (Tables 1-3). In contrast to the model without response change point (Fig. 3b), the change point model captures the variability of the entire *ε*_met_ dataset (Fig. 3c). Thus, this integrated change point model corroborates the following findings. First, *ε*_met_ exhibits a response change point. Below this change point, *ε*_met_ is largely constant with values of ≈96‰. Above the change point, *ε*_met_ variability is high. Second, change point crossing may require long-term drought beyond a critical level. Third, March to July precipitation and annual atmospheric CO_2_ concentration may govern *ε*_met_ variability on short timescales when long-term drought is beyond the critical level. Note, all relationships indicated by modelling require experimental confirmation in controlled settings.

### Variability in *δ*D_g_ is predominantly controlled by fractionating metabolic processes

Metabolic fractionations in *δ*D_1_ and *δ*D_2_ have a strong weight on whole-molecule D variability, *δ*D_g_. Variance partitioning shows that *δ*D_1_ and *δ*D_2_ together account for 71% of the variance in *δ*D_g_ (Fig. 2d). By contrast, *δ*D_3_ to *δ*D_6R_ each account for 5.8% on average. Assuming the variability in *δ*D_3_ to *δ*D_6R_ reflects the combined influence of non-metabolic fractionation processes affecting all *δ*D_*i*_, such as leaf water D enrichment, metabolic fractionations in *δ*D_1_ and *δ*D_2_ together account for 59.4% of the variance in *δ*D_g_ (71%-2*5.8%). When years affected by metabolic fractionation are considered exclusively (green dots in Fig. 2c), metabolic fractionations in *δ*D_1_ and *δ*D_2_ together account for 74.2% of the variance in *δ*D_g_ (Notes S5). After excluding the period affected by metabolic fractionation from the analysis (1983 to 1995), all *δ*D_*i*_ exhibit similar degrees of variance and contribute similarly to *δ*D_g_ (Notes S6) consistent with expectations related to non-metabolic fractionation processes. In conclusion, here, *δ*D_g_ variability is predominantly controlled by fractionating metabolic processes unaccounted for in current models. Note, our estimations implicitly assume invariability of metabolic fractionations in *δ*D_3_ to *δ*D_6R_. This simplifies actual conditions and results in underestimation of the influence of metabolic fractionations on *δ*D_g_.

### Altered metabolism is associated with below-average yet not exceptionally low growth

Metabolic changes as reflected by changes in metabolic fractionation may impact on growth. To test this, we investigated the relationship between *ε*_met_ and tree-ring width. We found that when the fractionating metabolic processes were upregulated, significantly more narrow tree rings were formed compared to when they were downregulated (Fig. 3d, one-tailed t-test: green vs. black circles, 0.75 vs. 0.99 mm, *p*<.05, *n*=9 and 22). Thus, altered metabolism is associated with below-average growth. However, equally narrow tree rings were formed in >50% of the years in which the fractionating metabolic processes were downregulated (Fig. 3d; tree-ring widths <0.99 mm, *n*=12). Thus, upregulations of the fractionating metabolic processes are not associated with exceptional growth declines.

## Discussion

### *ε*_met_: An isotope biomarker reports metabolic changes and drought

Here, we found highly variable metabolic fractionation signals at glucose H^1^ and H^2^ (Figs. 1-2). We approximated these signals by calculating *ε*_met_ (Eq. 5) and propose *ε*_met_ is a sensitive isotope biomarker reflecting changes in plant metabolism. Furthermore, we found a change point in *ε*_met_ (Step 1, Figs. 3a-c) suggesting *ε*_met_ features a two-state property. This property promises unambiguous identification of alternate metabolic states; a stable state where the fractionating metabolic processes remain downregulated, and a state where these processes are upregulated (upon crossing a change point). By contrast, other biomarkers develop in a purely linear manner with indistinct transitions between states hindering attempts to ascertain plant functioning. While the mechanisms behind other biomarkers in plant archives often remain elusive, *ε*_met_ can be (i) linked to specific physiological processes and (ii) incorporated into fractionation models (see below). Upon process elucidation, *ε*_met_ may help to find plants with the makeup to support crucial ecosystem services such as food and resource security.

As indicated by modelling, atmospheric CO_2_ concentrations and drought exert control over *ε*_met_ in *Pinus nigra* studied here (Tables 1-3, Figs. 3a-c). Values of *ε*_met_ >105‰ indicate long-term drought events affecting plant metabolism (Figs. 3a-c). Given careful sample selection (e.g., designed to avoid bias related to tree age), this may enable identification of historical occurrences. Fitting a model to *ε*_met_ values >105‰ (Fig. 3b, *ε*_met_=1802-0.566*PRE*-4.185*C*_a_) may enable analyses of metabolic effects of short-term drought and *C*_a_ during long-term drought events (by interpretation of the respective model terms), and reconstructions of short-term drought events during long-term drought events (solve for *PRE*, historical *C*_a_ data available).

*Pinus nigra* can access water from upper soil layers as well as from deep water sources (Stokes *et al.*, 2002; Peñuelas & Filella, 2003). Accordingly, at our study site, metabolic responses apparently require long-term drought (possibly involving groundwater depletion). In the absence of long-term drought, *ε*_met_ is blind to variability in short-term drought (possibly involving low soil moisture) and *C*_a_ (Figs. 3a-c). By contrast, species without groundwater access may provide this information. Lastly, *ε*_met_ may not always report drought but it may always report source-limited growth conditions (*cf.* Wieloch *et al.*, 2021a).

### The role of leaf intercellular CO_2_ concentrations

As indicated by modelling, upregulations of the fractionating metabolic processes occur in response to drought, yet high atmospheric CO_2_ concentrations exert counteracting effects (Tables 1-3, Figs. 3a-c). Isohydric plant species such as *Pinus nigra* respond to drought by closing their stomata (Sade *et al.*, 2012). This impedes CO_2_ uptake and promotes low intercellular CO_2_ concentrations, *C*_i_. By contrast, increasing *C*_a_ promotes increasing *C*_i_. Thus, upregulations of the fractionating metabolic processes may be mediated by low *C*_i_. In principle, whole-molecule stable carbon isotope ratios of plant organic matter, *δ*^13^C, enable *C*_i_ estimations because ^13^C fractionation during carbon uptake is related to *C*_i_/*C*_a_ (Farquhar *et al.*, 1982; Evans *et al.*, 1986). However, in the samples studied here, processes that are independent of carbon uptake fractionation control *δ*^13^C at three out of six glucose carbon positions (Wieloch *et al.*, 2018, 2021c,d). Therefore, we decided against using *δ*^13^C-derived *C*_i_ estimates to test this hypothesis.

### Stable isotope methodology

Variability in *δ*D_g_ is predominantly controlled by metabolic fractionation at glucose H^1^ and H^2^ (Fig. 2d). Since *δ*D_g_ can be measured by high-throughput isotope ratio mass spectrometry, a technique accessible to numerous laboratories, we investigated possibilities to (i) detect *δ*D_g_ datasets affected by metabolic fractionation, (ii) separate *δ*D_g_ datapoints affected by metabolic fractionation from other datapoints, and (iii) retrieve information from *δ*D_g_ about metabolic fractionation (Notes S7).

We found our *δ*D_g_ dataset holds clues to effects by metabolic fractionation at glucose H^1^ and H^2^ (Notes S7). Furthermore, separation of *δ*D_g_ datapoints affected by metabolic fractionation from other datapoints seems feasible yet not with high confidence. Discarding datasets/datapoints affected by metabolic fractionation at glucose H^1^ and H^2^ may improve *δ*D_g_ analyses of non-metabolic fractionation processes. However, analytical results will be impaired by other metabolic fractionations.

Modelling *ε*_met_, we found both a response change point marking the onset of a period with conditions favourable for upregulation of metabolic fractionation processes and environmental dependences of upregulation (Table 3, Eq. 7, Fig. 3c). Applying this model to *δ*D_g_, we found the same change point, but the environmental dependences of upregulation were not sufficiently constrained for interpretation (Notes S7). Thus, by itself, *δ*D_g_ analysis of glucose does not yield robust information about the environmental dependences of metabolic fractionation at H^1^ and H^2^. However, in combination with modelled or measured leaf water *δ*D values, information on overall metabolic fractionation may be retrieved (*cf.* Cormier *et al.*, 2018). By contrast, resolving information about different metabolic fractionations requires the intramolecular approach.

Current *δ*D_g_ models describe metabolic fractionations as being constant (Roden *et al.*, 2000). As shown here, this is inadequate for plants grown under dry conditions where metabolic fractionations can exert predominant control over *δ*D_g_ variability (Fig. 2d). Thus, to model plant D abundances over the whole range of environmental conditions, metabolic fractionations need to be incorporated as variables which requires detailed knowledge about underlying processes.

Metabolic fractionation processes known to affect tree-ring glucose H^1^ and H^2^ cannot explain our findings. Specifically, *Picea abies* reportedly exhibits similar heterotrophic hydrogen exchange rates at tree-ring glucose H^1^ and H^2^ (≈42%) (Augusti *et al.*, 2006). Since *Pinus nigra* as a close relative can be expected to behave similarly, heterotrophic hydrogen exchange at these positions cannot cause pronounced deviations from slope=1 as observed for the *δ*D_1_-*δ*D_2_ regression (Fig. 2c). Therefore, we will now derive experimentally testable theories on the metabolic origins of the fractionation signals reported here.

### Theory 1. Isotope fractionation related to sucrose-to-starch carbon partitioning

We found a highly variable metabolic fractionation signal at tree-ring glucose H^2^ (*ε*_met_ at H^2^: *SD*≈±79‰, *range*≈260‰, *n*=31) which occurs upon the crossing of a response change point (Figs. 3a-c). At the leaf level, tree-ring glucose has two precursors, starch and sucrose. Growing *Phaseolus vulgaris* and *Spinacia oleracea* under optimal conditions, Schleucher *et al.* (1999) reported D depletions at glucose H^2^ of leaf starch relative to sucrose of 333‰ and 500‰, respectively. This difference may contribute to the signal at tree-ring glucose H^2^ when the relative flux of assimilated carbon into starch and sucrose changes (Fig. 4). Sucrose-to-starch carbon partitioning ratios are controlled primarily by the rate of carbon assimilation (Sharkey *et al.*, 1985) and were shown to increase with decreasing assimilation rates (Sharkey *et al.*, 1985), decreasing *C*_i_ (Sharkey *et al.*, 1985), decreasing light (Sharkey *et al.*, 1985; Quick *et al.*, 1989), and increasing drought (Quick *et al.*, 1989, 1992; Vassey & Sharkey, 1989). Reported responses are functionally coherent and qualitatively consistent across all species studied. Furthermore, responses of sucrose-to-starch carbon partitioning to ecophysiological changes exhibit change points. For instance, in leaves of *Phaseolus vulgaris*, the sucrose-to-starch carbon partitioning ratio was found to be largely constant at *C*_i_≥150 ppm yet increases below this change point from ≈0.5 to ≈4 (Sharkey *et al.*, 1985). Thus, as *C*_i_ drops below the change point, the relative contribution of leaf sucrose to tree-ring glucose biosynthesis may increase, and tree-ring glucose H^2^ may gradually approach the higher relative D abundance of leaf sucrose.

**Figure 4:**
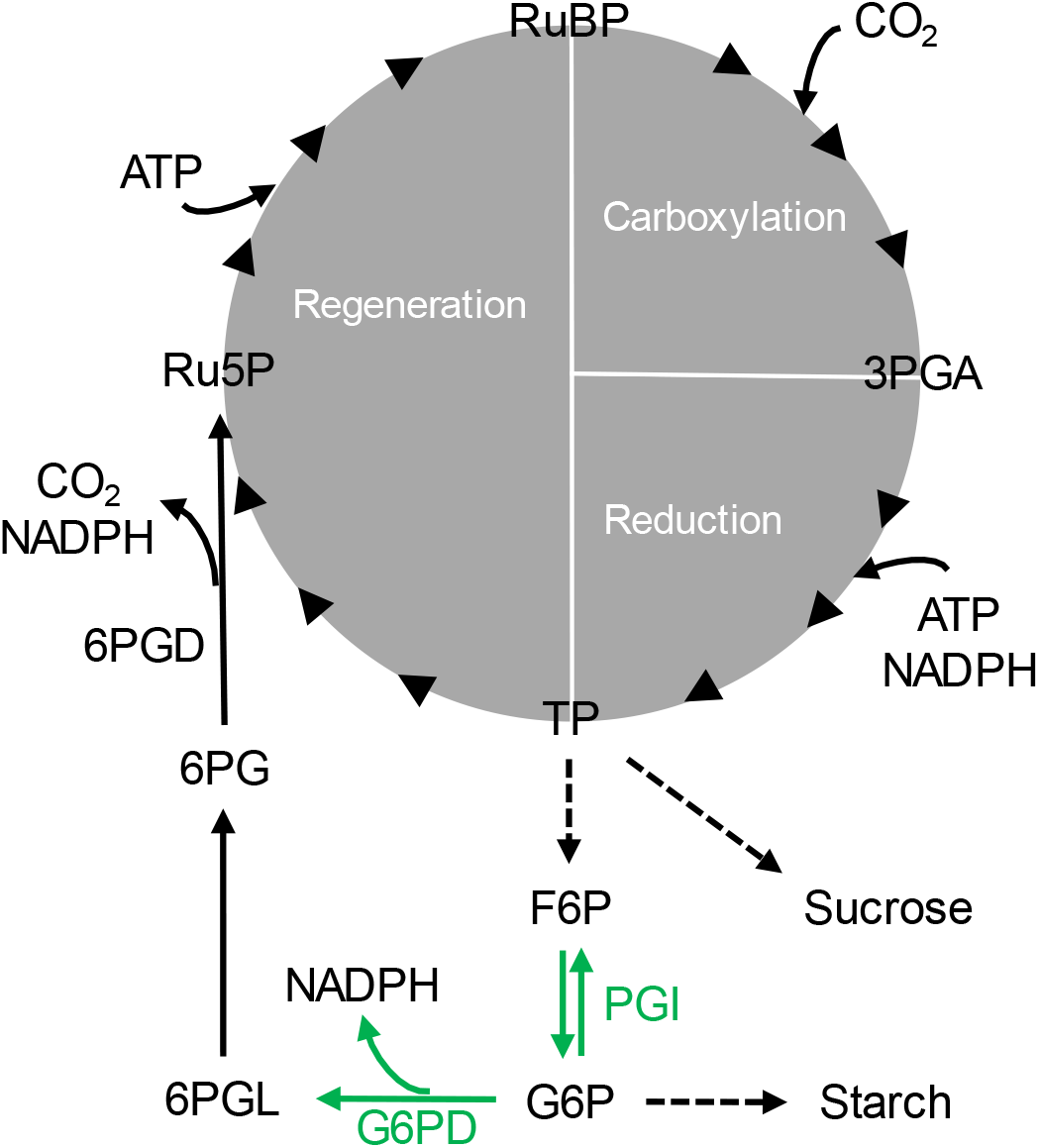
Sucrose-to-starch carbon partitioning and anaplerotic carbon flux into the Calvin-Benson cycle. Green: Enzyme reactions proposedly introducing D isotope signals at glucose H^1^ and H^2^. G6PD has a kinetic isotope effect of *α*_D_=k_H_/k_D_=2.97 (Hermes *et al.*, 1982) while PGI has a kinetic isotope effect of 2.22 in the F6P to G6P direction and an equilibrium isotope effect of 1.11 in G6P (Rose & O’Connell, 1961; Wieloch *et al.*, 2021a). Dashed arrows: Intermediate reactions not shown. Enzymes: 6PGD, 6-phosphogluconate dehydrogenase; G6PD, glucose-6-phosphate dehydrogenase; PGI, phosphoglucose isomerase. Metabolites: 3PGA, 3-phosphoglycerate; 6PG, 6-phosphogluconate; 6PGL, 6-phosphogluconolactone; ATP, adenosine triphosphate; F6P, fructose 6-phosphate; G6P, glucose 6-phosphate; NADPH, nicotinamide adenine dinucleotide phosphate; Ru5P, ribulose 5-phosphate; RuBP, ribulose 1,5-bisphosphate; TP, triose phosphates (glyceraldehyde 3-phosphate, dihydroxyacetone phosphate).

### Theory 2. Isotope fractionation related to anaplerotic carbon flux into the Calvin-Benson cycle

Metabolic fractionation signals at tree-ring glucose H^1^ and H^2^ are closely related (Figs. 2a, and c). We propose this is due to (i) isotope fractionation related to anaplerotic carbon flux into the Calvin-Benson cycle (CBC) affecting both positions, and (ii) dependences of both anaplerotic carbon flux into the CBC and sucrose-to-starch carbon partitioning on the rate of carbon assimilation.

Sharkey and Weise (2016) proposed the CBC is anaplerotically refilled by injection of five-carbon sugar phosphates from the stromal pentose phosphate pathway with glucose 6-phosphate (G6P) as precursor (Fig. 4). Primary flux control is believed to be exerted at the level of stromal phosphoglucose isomerase (PGI) which catalyses interconversions between fructose 6-phosphate (F6P) and G6P. Under optimal growth conditions, the PGI reaction is strongly removed from equilibrium on the side of F6P resulting in low stromal G6P concentrations (Dietz, 1985; Gerhardt *et al.*, 1987; Kruckeberg *et al.*, 1989; Schleucher *et al.*, 1999). Proposedly, low G6P concentrations restrict the anaplerotic flux (Sharkey & Weise, 2016). However, as the PGI reaction shifts towards equilibrium and G6P synthesis, anaplerotic flux into the CBC is believed to increase (Sharkey & Weise, 2016). This involves stromal glucose-6-phosphate dehydrogenase (G6PD), the first enzyme of the pentose phosphate pathway. In the light, G6PD is downregulated via redox regulation by thioredoxin (Née *et al.*, 2009). However, downregulation can be allosterically reversed by increasing G6P concentrations (Cossar *et al.*, 1984; Preiser *et al.*, 2019).

*Phaseolus vulgaris* and *Spinacia oleracea* grown under optimal conditions exhibit pronounced D depletions at glucose H^2^ of leaf starch relative to sucrose (see above) (Schleucher *et al.*, 1999). These depletions were attributed to the stromal PGI reaction being more strongly removed from equilibrium ([G6P]/[F6P]=3.31) than the cytosolic PGI reaction (Dyson & Noltmann, 1968; Schleucher *et al.*, 1999). Thus, as the stromal PGI reaction shifts towards equilibrium, H^2^ of stromal G6P and its derivatives including leaf starch and tree-ring glucose may become less D depleted. Furthermore, G6PD catalyses the irreversible conversion of G6P to 6-phosphogluconolactone and was shown to have a strong D isotope effect i*n vitro* (*α*=2.97) (Hermes *et al.*, 1982). Thus, upregulation of the anaplerotic flux may cause D enrichments at H^1^ of stromal G6P and its derivatives.

As indicated by modelling, glucose H^1^ and H^2^ become D enriched as drought increases and *C*_a_ decreases beyond a change point (Tables 1-3, Figs. 3a-c). Increasing drought and decreasing *C*_a_ may cause decreasing *C*_i_ (see above) which may result in decreasing carbon assimilation. As carbon assimilation decreases below a change point, the stromal PGI reaction shifts towards equilibrium and increased G6P concentrations (Dietz, 1985). This may cause increasing G6PD activity and anaplerotic flux. Thus, expanding theory by Sharkey and Weise (2016), we propose anaplerotic flux into the CBC is upregulated in response to decreasing carbon assimilation, *C*_i_, and *C*_a_, and increasing drought beyond a change point.

Analysis of intramolecular ^13^C/^12^C ratios of the samples studied here has already provided evidence consistent with anaplerotic flux into the CBC at high VPD (Wieloch *et al.*, 2018). These authors explained a strong common ^13^C signal at tree-ring glucose C-1 and C-2 by anaplerotic flux changes. For shifts of the PGI reaction towards equilibrium, they predicted ^13^C/^12^C increases at C-1 and C-2 at a ratio of 2.25 and found a ratio of 2.74 (+1.35SE, −0.60SE) confirming their prediction. However, changes in sucrose-to-starch carbon partitioning would have the same effect. Furthermore, two recent studies report isotope and gas exchange evidence consistent with anaplerotic flux in *Helianthus annuus* at low *C*_i_ (Wieloch *et al.*, 2021a,b). That said, other mechanisms than those discussed here may contribute to the signals at tree-ring glucose H^1^ and H^2^. This includes carbon flux into the oxidative branch of the cytosolic pentose phosphate pathway, shifts of the cytosolic PGI reaction, and secondary isotope effects both in leaf and tree-ring cells.

### Metabolic fractionation signals at H^1^ and H^2^: A general phenomenon in C_3_ plants?

Cormier *et al.* (2018) grew six phylogenetically diverse angiosperms under varying *C*_a_. For leaf α-cellulose, these authors reported average whole-molecule *ε*_met_ increases of ≈20‰ in response to *C*_a_ decreases from 280 to 150 ppm yet no significant change above 280 ppm. Hence, their observations are qualitatively in line with ours (Tables 1-3, Figs. 3a-c). However, our data show an average *ε*_met_ increase at tree-ring glucose H^1^ and H^2^ of up to 150‰ (Fig. 2b). Assuming ≈40% heterotrophic hydrogen exchanges as observed at these positions in *Picea abies* (Augusti *et al.*, 2006), we calculate an undiluted leaf-level increase of ≈250‰ (150‰/60*100) which corresponds to a whole-molecule increase of ≈70‰ (2*250‰/7). Thus, *ε*_met_ increases in the gymnosperm *Pinus nigra* are considerably larger than the ≈20‰ angiosperm average effect reported by Cormier *et al.* (2018). Additionally, whole-molecule data have a limited interpretability (*cf. ‘*Introduction’) and effects reported by Cormier *et al.* (2018) may be located at either of the seven H-C positions within α-cellulose glucose. Thus, it remains uncertain whether the fractionation signals reported here occur generally in C_3_ plants.

### Concluding remarks

Stable isotope analysis on plant archives such as tree rings may convey information (*inter alia*) about plant acclimation and adaptation (e.g., to CO_2_ fertilisation), biosphere-atmosphere CO_2_ exchange, and paleoclimate trends. This long-term perspective is inaccessible to manipulation and monitoring experiments yet crucial for understanding plant and Earth system functioning. The better we understand plant isotope fractionation, the more information can be retrieved from archives (at higher quality). The present paper and recent studies on metabolic fractionation (Wieloch *et al.*, 2018, 2021c, a,d; Ladd *et al.*, 2021) show there is much room for improvement. Thus, we recommend pushing the development of intramolecular isotope methodology.

## Acknowledgements

For helpful discussions, we thank Thomas Cech (Austrian Research Centre for Forests), Youyi Fong (Fred Hutchinson Cancer Research Center, Seattle), Erhard Halmschlager (University of Natural Resources and Life Sciences, Vienna), John D. Marshall (SLU Umeå), John S. Roden (Southern Oregon University), Thomas D. Sharkey (Michigan State University), and Roland A. Werner (ETH Zürich). The research of I.E., and J.S. and part of the research of T.W. was supported by the Swedish Research Council VR (2013-05219, 2018-04456), the Knut and Alice Wallenberg Foundation (“NMR for Life” facility and grant 2015.0047) and the Kempe foundations.

## Author contributions

T.W., and J.S. conceived the study. T.W. led the research. T.W., M.G., H.S., J.S., and I.E. collected and prepared the samples and acquired data. T.W., and J.Y. analysed the data. T.W. developed theories about the origin of reported isotope signals. T.W. wrote the paper with input from A.A., H.S., J.S., and J.Y.

## Data availability

The data supporting the findings of this study are available from the corresponding author upon reasonable request.

## Supporting information

Notes S1. Statistical analyses

Notes S2. Grouping of annual *δ*D_*i*_ patterns of tree-ring glucose by HCA

Notes S3. Histograms of *ε*_met_

Notes S4. Bivariate relationships of *ε*_met_ with precipitation and atmospheric CO_2_ concentration

Notes S5. Contributions of *δ*D_*i*_ to the variance in *δ*D_g_ after excluding data not affected by the fractionating metabolic processes

Notes S6. Contributions of *δ*D_*i*_ to the variance in *δ*D_g_ after excluding data affected by the fractionating metabolic processes

Notes S7. Metabolic fractionation at the whole-molecule level

