## Supporting information for "Metabolism is a major driver of hydrogen isotope fractionation recorded in tree-ring glucose of *Pinus nigra*"

**Notes S1. Statistical analyses**

Hierarchical cluster analysis was performed using the R function hclust() of the STATS package on z-scores of $\delta D_i$ using Euclidean distances and Ward's fusion criterion for cluster formation (n=7*31). Mood's median tests were performed using the R function median_test() of the COIN package. Batch change point analysis based on the non-parametric Mann-Whitney-Wilcoxon test was performed using the R function detectChangePointBatch() of the CPM package (Ross, 2015). Multiple linear regression modelling was performed using the R function lm() of the STATS package. Change point regression modelling (step type) was performed using the R function chngptm() of the CHNGPT package (Fong *et al.*, 2017). Statistical significances of the change point and the change point model were calculated using the R functions chngpt.test() of the CHNGPT package and lrtest() of the LMTEST package, respectively.

**Notes S2. Grouping of annual $\delta D_i$ patterns of tree-ring glucose by HCA**

Metabolic fractionations affect specific intramolecular H-C positions and thus introduce intramolecular $\delta D_i$ patterns (Figs. 1, 2b, and S2b). To identify annual patterns that were similarly/differently modified by metabolic fractionation, we performed a Hierarchical Cluster Analysis (HCA) on annual $\delta D_i$ patterns (Fig. S2a). We found two groups of $\delta D_i$ patterns: a high-value and a low-value group according to $\delta D_1$ and $\delta D_2$ values (Figs. 2b, and S2b). Please note that the population to which our data belong must be continuously distributed despite the apparent grouping and the gap in $\delta D_2$ in the ≈200 to 250‰ range (Figs. S2a, and S2b). This is because tree rings record ecophysiological information continuously over the course of growing seasons, i.e., the temporal impact of metabolic fractionations at $H^1$ and $H^2$ principally varies in a continuous way. Nevertheless, the apparent grouping in our data is convenient to investigate causes of high and low $\delta D_1$ and $\delta D_2$ values. Please note that if our dataset had been continuously distributed, we could still have arbitrarily separated it into low- and high-value groups and investigated underlying causes. However, in the present case, HCA separates the data for us.

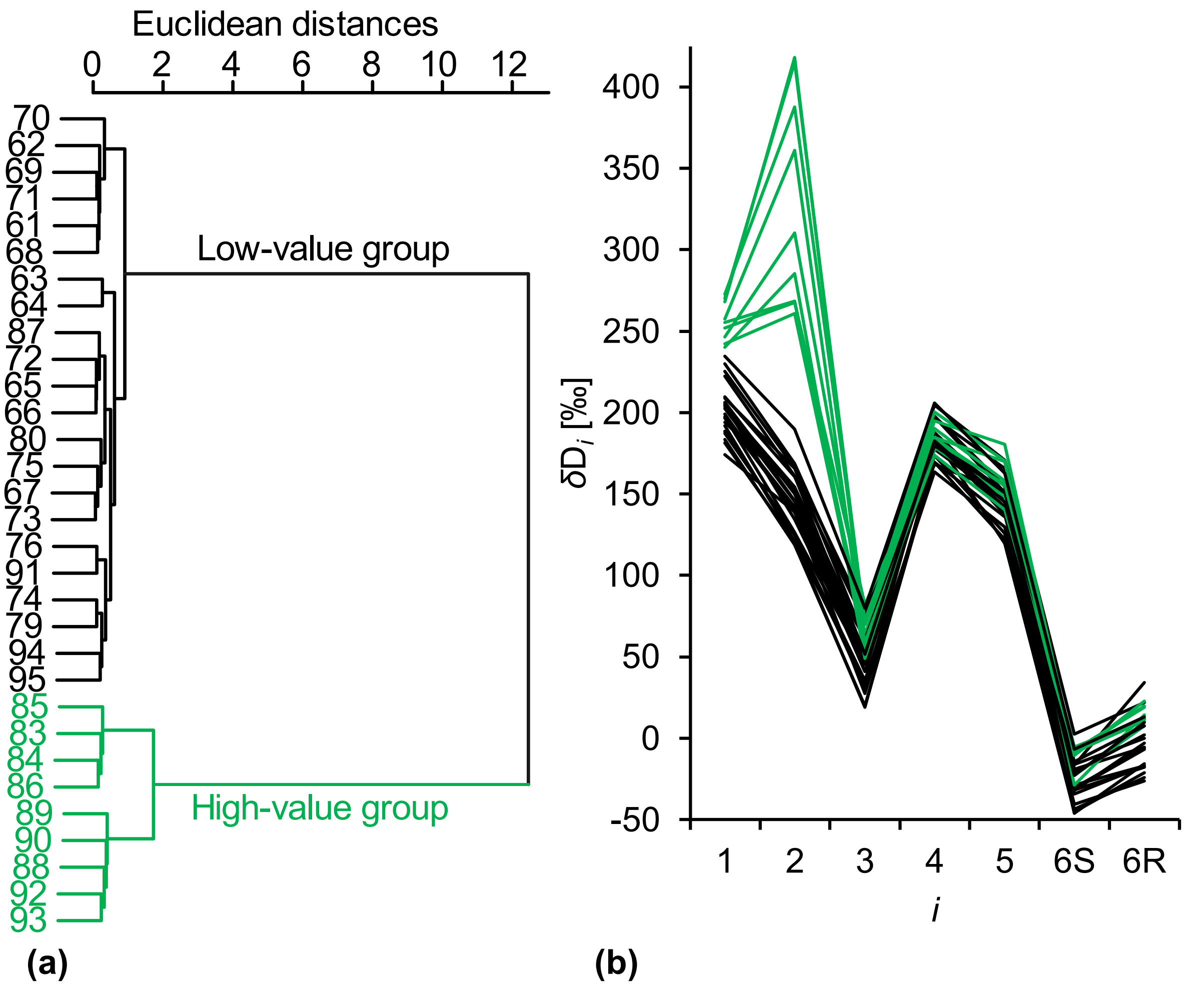


**Figure S2 (a)** Grouping of annual $\delta D_i$ patterns by Hierarchical Cluster Analysis (HCA). **(b)** Annual $\delta D_i$ patterns. $\delta D_i$ denotes D abundances at intramolecular H-C positions in tree-ring glucose. Data were acquired for tree-ring glucose of *Pinus nigra* laid down from 1961 to 1995 at a site in the Vienna basin (±*SE*=5.4‰, $n \geq 3$). Prior to HCA, outliers were replaced by timeseries averages. Data reference: Average D abundance of the methyl-group hydrogens of the glucose derivative used for NMR measurements. Figure S2b shows discrete data. Lines used to guide the eye.

### Notes S3. Histograms of $\varepsilon_{met}$

Figures S3a, and b show histograms of $\varepsilon_{met}$ including all data, 1961 to 1995, and data of 1983 to 1995, respectively. The dataset including all data clearly does not follow a normal distribution. By contrast, there are no indications that the dataset including only years in which groundwater levels were low deviates from a normal distribution (1983 to 1995, Fig. 3a).

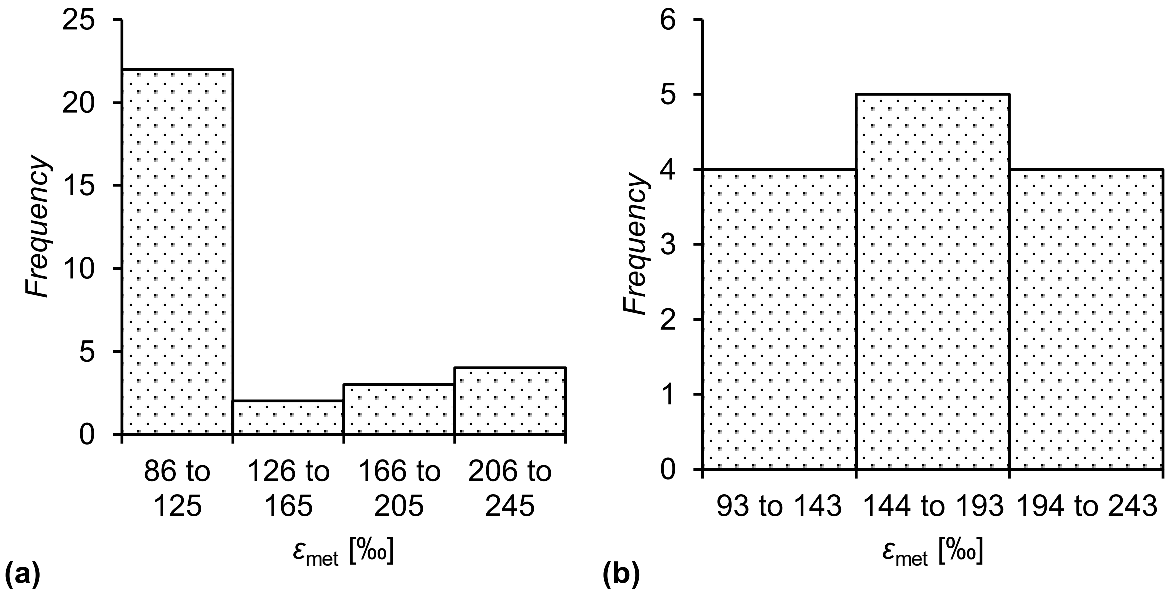


**Figure S3 (a)** Histogram of average metabolic fractionation at glucose $H^1$ and $H^2$, $\varepsilon_{met}$, including all data, 1961 to 1995 (*n*=31). **(b)** Histogram of $\varepsilon_{met}$ including data of 1983 to 1995 corresponding to years with low groundwater level (*n*=13, Fig. 3a). Data for the calculation of $\varepsilon_{met}$ were acquired for tree-ring glucose of *Pinus nigra* laid down from 1961 to 1995 at a site in the Vienna basin (±*SE*=3.5‰, $n \geq 3$).

**Notes S4. Bivariate relationships of $\varepsilon_{met}$ with precipitation and atmospheric $CO_2$ concentration**

Figures S4a, and b show bivariate relationships of $\varepsilon_{met}$ with March to July precipitation, *PRE*, and annual atmospheric $CO_2$ concentrations, $C_a$, respectively, for two groups of data. While the first group includes data of 1961 to 1982 corresponding to high groundwater storage (black), the second group includes data of 1983 to 1995 corresponding to low groundwater storage (blue, Fig. 3a). Linear modelling shows that relationships of $\varepsilon_{met}$ with both environmental parameters are group specific. While the first group shows no significant relationships ($\varepsilon_{met}$ ~ *PRE*: $R^2$=0.21, $p$>.05, $n$=18; $\varepsilon_{met}$ ~ $C_a$: $R^2$=0.01, $p$>.7, $n$=18), the second group shows significant negative relationships ($\varepsilon_{met}$ ~ *PRE*: $R^2$=0.71, $p$<0.001, $n$=13; $\varepsilon_{met}$ ~ $C_a$: $R^2$=0.54, $p$<.01, $n$=13). Note that low $\varepsilon_{met}$ values can occur under low groundwater storage (Fig. 3b, 1987, 1991, 1994, and 1995). This is explained by high precipitation during spring and summer and high atmospheric $CO_2$ concentrations (Figs. S4a, and b, filled circles).

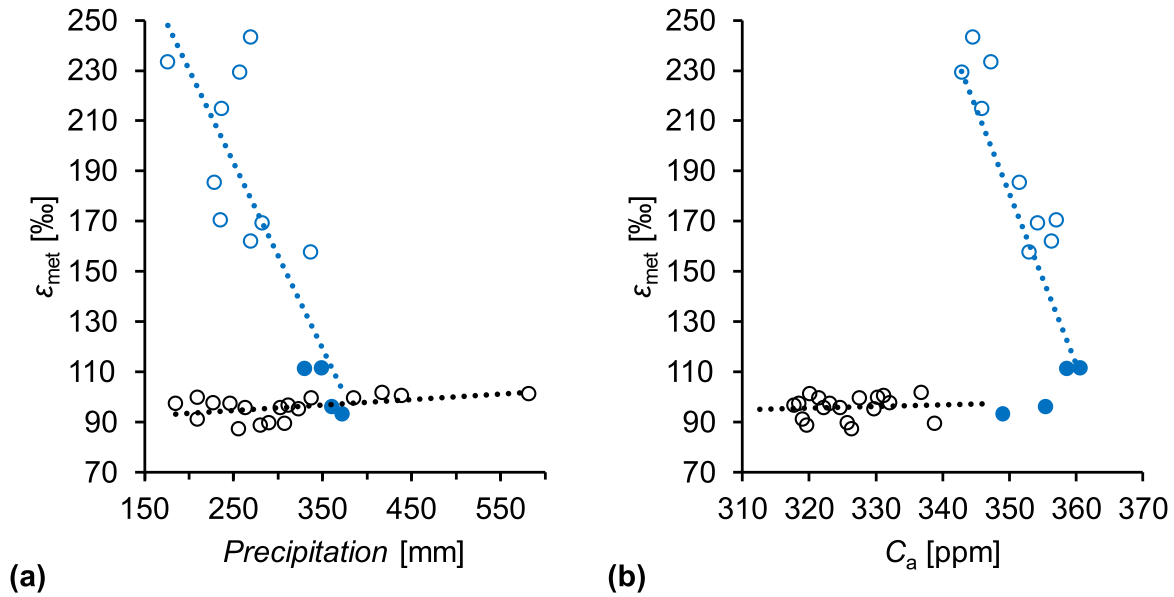


**Figure S4 (a)** Relationship between average metabolic fractionation at glucose $H^1$ and $H^2$, $\varepsilon_{met}$, and March to July precipitation. **(b)** Relationship between $\varepsilon_{met}$ and annual atmospheric $CO_2$ concentration, $C_a$. Black and blue circles: Data of years with high groundwater storage, 1961 to 1982, and low groundwater storage, 1983 to 1995, respectively (Fig. 3a). Filled circles: Data of 1987, 1991, 1994, and 1995. Dotted black and blue lines: Trendlines pertaining to datasets of the same colour. Data for the calculation of $\varepsilon_{met}$ were acquired for tree-ring glucose of *Pinus nigra* laid down from 1961 to 1995 at a site in the Vienna basin (±*SE*=3.5‰, *n*≥3).

### Notes S5. Contributions of $\delta D_i$ to the variance in $\delta D_g$ after excluding data not affected by the fractionating metabolic processes

Metabolic fractionations in $\delta D_1$ and $\delta D_2$ have a strong weight on whole-molecule D variability, $\delta D_g$. To assess this weight exclusively for years with upregulated fractionating metabolic processes, we repeated the variance partitioning on corresponding data but exclude 1988 and 1990 because of data gaps as result of the outlier analysis (1983-1986, 1989, 1992, and 1993; Fig. 2c). We found that $\delta D_1$ and $\delta D_2$ together account for 86.8% of the variance in $\delta D_g$. By contrast, $\delta D_3$ to $\delta D_5$ each account for 6.3% on average. Interestingly, $\delta D_{6S}$ and $\delta D_{6R}$ reduce the variability of $\delta D_g$ by -2.8% on average. Assuming the variability in $\delta D_3$ to $\delta D_5$ reflects the combined influence of known fractionation processes affecting all $\delta D_i$, such as leaf water D enrichment, metabolic fractionations in $\delta D_1$ and $\delta D_2$ together account for 74.2% of the variance in $\delta D_g$ (86.8%-2*6.3%).

**Notes S6. Contributions of $\delta D_i$ to the variance in $\delta D_g$ after excluding data affected by the fractionating metabolic processes**

Metabolic processes have strong effects on $\delta D_1$, $\delta D_2$, and $\delta D_g$ (Figs. 1, and 2). After excluding years affected by these processes from the variance partitioning analysis (1983 to 1995), all $\delta D_i$ exhibit similar degrees of variance and contribute similarly to $\delta D_g$ (Fig. S6).

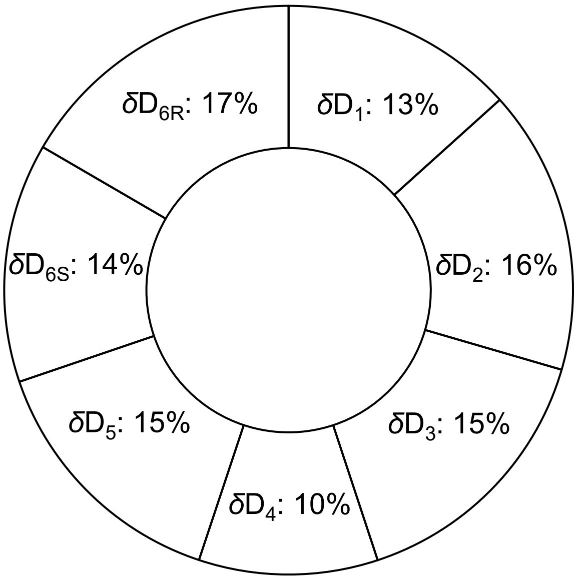


**Figure S6** Percentage contributions of $\delta D_i$ to the variance of $\delta D_g$. $\delta D_i$ and $\delta D_g$ denote timeseries of D abundances at intramolecular H-C positions in glucose and of the whole molecule, respectively. Data were acquired for tree-ring glucose of *Pinus nigra* laid down from 1961 to 1982 at a site in the Vienna basin ($\delta D_i$: $\pm SE$=5.5‰, $n \geq 3$; $\delta D_g$: $\pm SE$=3.7‰, $n \geq 3$). Outliers were removed prior to analysis. The analysis is based on years without missing data ($n$=8*14). Data reference: Average D abundance of the methyl-group hydrogens of the glucose derivative used for NMR measurements.

**Notes S7. Metabolic fractionation at the whole-molecule level**

Within this paragraph, the term ‘metabolic fractionation’ refers to metabolic fractionation at glucose $H^1$ and $H^2$.

Variability in $\delta D_g$ is predominantly controlled by metabolic fractionation (Fig. 2d). Since $\delta D_g$ can be measured by high-throughput isotope ratio mass spectrometry (a technique accessible to numerous laboratories), we will now investigate possibilities to (i) identify $\delta D_g$ datasets affected by metabolic fractionation, (ii) separate $\delta D_g$ datapoints affected by metabolic fractionation from other datapoints, and (iii) retrieve information from $\delta D_g$ about metabolic fractionation.

(i) Metabolic fractionation caused occasional $\delta D_g$ increases above normal $\delta D_g$ values but never $\delta D_g$ decreases (green dots in Fig. S7a). Consequently, the $\delta D_g$ distribution is asymmetrical with a moderate positive skew, has increased variability, and is nearly significantly different from normality (Fig. S7c; *skewness*=0.55, *range*=83.2‰, *SD*=23.8‰; Shapiro-Wilk normality test: $W$=0.93616, $p$=0.12). After excluding data affected by metabolic fractionation, the $\delta D_g$ distribution is approximately symmetrical, has lower variability, and is not significantly different from normality (Fig. S7d; *skewness*=0.32, *range*=47.6‰, *SD*=13.5‰; Shapiro-Wilk normality test: $W$=0.95866, $p$=0.58). Furthermore, we found a change point in the complete $\delta D_g$ timeseries (non-parametric Mann-Whitney-Wilcoxon test: $p$<.001, $n$=25) which corresponds to the change point in $\varepsilon_{met}$ and marks the onset of a period with conditions favourable for upregulations of the fractionating metabolic processes (main text, '*Step 1*'). Thus, both visual inspection of the $\delta D_g$ distribution and statistical tests indicate effects by metabolic fractionation.

Our $\delta D_g$ dataset is relatively small ($n$=25) and, therefore, not an ideal approximation of the underlying probability distribution. Theoretically, we would expect a bimodal distribution. Data not affected by metabolic fractionation (black dots in Fig. S7a) would be represented by a low-value peak in the histogram. Data affected by metabolic fractionation (green dots in Fig. S7a) would be represented by a high-value peak adjacent to the low-value peak. The relative height of these peaks would depend on the relative frequency of long-term drought events (groundwater depletions below the critical level).

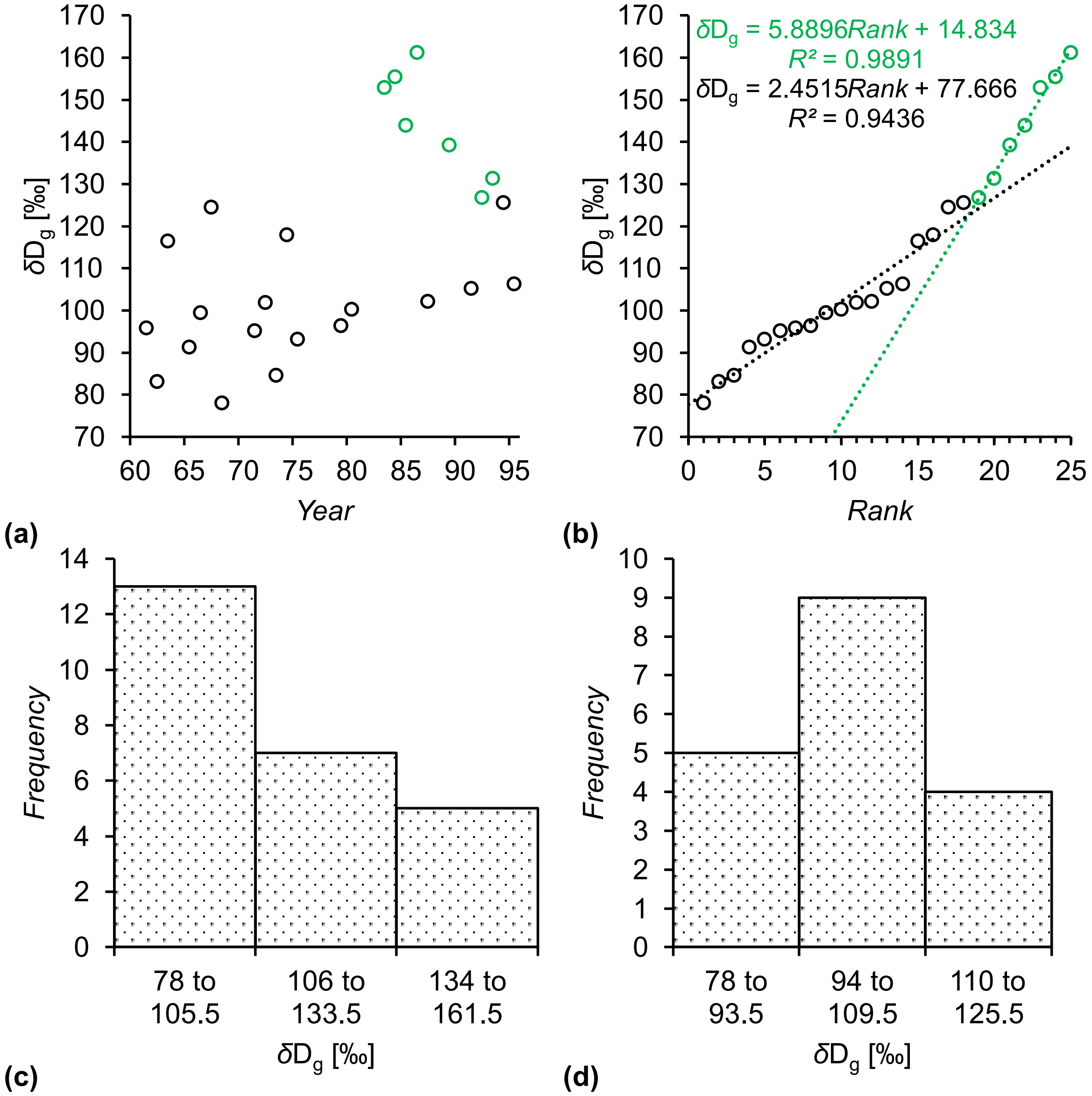


**Figure S7 (a)** Timeseries of whole-molecule deuterium abundance, $\delta D_g$. Green and black dots, data affected and not affected by metabolic fractionation at glucose $H^1$ and $H^2$, respectively ($n$=7, and $n$=18). **(b)** $\delta D_g$ ranked according to value from low to high. Green and black lines, trendlines pertaining to data affected and not affected by metabolic fractionation at glucose $H^1$ and $H^2$, respectively. **(c)** Histogram of $\delta D_g$ including all data ($n$=25, black and green dots in Fig. S7a). **(d)** Histogram of $\delta D_g$ excluding data affected by metabolic fractionations at glucose $H^1$ and $H^2$ ($n$=18, black dots in Fig. S7a). Data were acquired for tree-ring glucose of *Pinus nigra* laid down from 1961 to 1995 at a site in the Vienna basin (±*SE*=3.4‰, $n \geq 3$). Outliers were removed prior to analysis. Data reference: Average D abundance of the methyl-group hydrogens of the glucose derivative used for NMR measurements.

(ii) Figure S7a shows $\delta D_g$ as function of time with green dots representing data affected by metabolic fractionation (*cf.* Fig. 2c). Without colour coding, a clear separation between datapoints affected by metabolic fractionation and other datapoints is not feasible. Figure S7b shows the same data ranked by value from low to high. Data affected by metabolic fractionation have the highest ranks and are sitting neatly on a line (green line, $R^2$=0.99, $n$=7). The slope of this line is 2.4 times steeper than the slope of the line pertaining to data not affected by metabolic fractionation (black line, $R^2$=0.94, $n$=18). This may enable $\delta D_g$ data separation yet not with high confidence. For instance, without colour coding, it is unclear whether the four datapoints before the green datapoints were also affected by metabolic fractionation. Furthermore, if the number of data affected by metabolic fractionation was 2.4 times higher ($n$≈17), both lines would have the same slope.

(iii) A change point model explains most of the variance in $\varepsilon_{met}$ (main text, '*Model 2*', $R^2$=0.94, $p<10^{-15}$, $n$=31, Eq. 7), and all explanatory variables contribute significantly to this model (Table 3). While the same change point model explains a significant fraction of the variance in $\delta D_g$ ($R^2$=0.76, $p<10^{-5}$, $n$=25, Eq. 7), most explanatory variables do not contribute significantly (Table S7). Thus, at the level of $\delta D_g$, the environmental dependences of the fractionating metabolic processes are insufficiently constraint for interpretation.

**Table S7** Estimated coefficients of the $\delta D_g$ change point model.

| Coefficient | Estimate | SE* | Lower 95% CI | Upper 95% CI | p-value* |
|---|---|---|---|---|---|
| **$\beta_1$** | 638.12 | 328.99 | -246.67 | 1042.99 | 0.05 |
| **$\beta_2$** | -0.21463 | 0.09973 | -0.40477 | -0.01382 | <.05 |
| **$\beta_3$** | -1.2620 | 0.9331 | -2.4785 | 1.1794 | 0.2 |
| **$\beta_4$** | -562.80 | 730.68 | -2260.27 | 604.01 | 0.4 |
| **$\beta_5$** | 0.18721 | 0.13934 | -0.07207 | 0.47415 | 0.2 |
| **$\beta_6$** | 1.3598 | 2.3345 | -2.0441 | 7.1071 | 0.6 |
| **e** | -0.70310 | 0.23832 | -0.85880 | 0.07540 | <.001 |

A change point model was fitted to measured whole-molecule deuterium abundances, $\delta D_g$ ($R^2$=0.76, $p<10^{-5}$, $n$=25, Eq. 7) (Fong *et al.*, 2017). Data were acquired for tree-ring glucose of *Pinus nigra* laid down from 1961 to 1995 at a site in the Vienna basin (±*SE*=3.4‰, $n\geq3$). $\beta_1$ to $\beta_6$, and e denote model coefficients (Eq. 7). SE and CI denote the standard error and confidence interval, respectively. Asterisks mark estimations which assume that bootstrap sampling followed a normal distribution.
